# MCSPACE: inferring microbiome spatiotemporal dynamics from high-throughput co-localization data

**DOI:** 10.1101/2024.12.06.627244

**Authors:** Gurdip Uppal, Guillaume Urtecho, Miles Richardson, Isin Y. Comba, Jeongchan Lee, Thomas Moody, Harris H. Wang, Georg K. Gerber

## Abstract

Recent advances in high-throughput approaches for estimating co-localization of microbes, such as SAMPL-seq, allow characterization of the biogeography of the gut microbiome longitudinally and at unprecedented scale. However, these high-dimensional data are complex and have unique noise properties. To address these challenges, we developed MCSPACE, a probabilistic AI method that infers from microbiome co-localization data spatially coherent assemblages of taxa, their dynamics over time, and their responses to perturbations. To evaluate MCSPACE’s capabilities, we generated the largest longitudinal microbiome co-localization dataset to date, profiling spatial relationships of microbes in the guts of mice subjected to serial dietary perturbations over 76 days. Analyses of these data and an existing human longitudinal dataset demonstrated superior benchmarking performance of MCSPACE over existing methods, and moreover yielded insights into spatiotemporal structuring of the gut microbiome, including identifying temporally persistent and dynamic microbial assemblages in the human gut, and shifts in assemblages in the murine gut induced by specific dietary components. Our results highlight the utility of our method, which we make available to the community as an open-source software tool, for elucidating dynamics of microbiome biogeography and gaining insights into the role of spatial relationships in host-microbial ecosystem function.

## Introduction

The mammalian gut harbors a highly complex and dynamic microbial ecosystem that modulates host physiology and is important for maintaining health^1–7^. Trillions of microbial cells, comprised of hundreds of diverse taxa colonizing throughout the gut, co-exist in a highly heterogeneous environment. Spatial features of this environment, including regional differences in nutrient availabilities, cross-feeding interactions^8^, and attachment sites^9^, strongly influence where particular taxa colonize. These features have functional implications, with spatial organization of microbes in the gut facilitating maintenance of biodiversity^10^, microbe-microbe^11^ and host-microbe^12^ interactions, as well as the stability and plasticity of the gut ecosystem^13–15^. In addition, the microbiome is temporally dynamic, particularly in the context of environmental perturbations^9,15,16^, such as changes in the host diet. Simultaneous spatial and temporal characterization of the gut microbiome is thus important for fully understanding this ecosystem, with potential to provide new insights into key questions including the kinetics of microbial community assembly^11,17^ and the development and maintenance of robust and stable microbial interdependencies^7,13,14,18^, which impact host physiology and disease pathophysiology^15,16,18^.

Although spatiotemporal characterization of the microbiome has the potential to provide many exciting new insights, technical and practical challenges have historically limited the throughput of spatially-resolved methods. Initial fluorescence in situ hybridization (FISH)-based methods were limited by spectral diversity and could only target a small number of taxa at a time^19–21^. Advancements in FISH probe multiplexing technologies have enabled the simultaneous detection of more taxa^19,22,23^ and interesting recent approaches combining metabolic labeling with FISH^24^ can provide multi-modal spatially resolved data. However, the throughput of these methods remains limited, because individual probes for each target microbe^22,23^ must be developed with considerable experimental optimization required^25,26^. Sequencing-based approaches using laser microdissection to characterize the microbiome in regions of the gut^27,28^ do not require design of individual probes, and can thus assess the full array of microbes present, but are limited in spatial resolution and throughput due to the microdissection process. Spatial transcriptomics technologies offer another promising approach for studying microbiome biogeography^12,29^, but are at present relatively low throughput and very costly.

Recently introduced sequencing-based technologies for elucidating spatial co-localization of microbes, such as MaPS-seq^30^ and its successor SAMPL-seq^31^ that further improved scalability using combinatorial split-and-pool barcoding^32–34^, overcame many of the above-mentioned throughput challenges and are thus attractive methods for spatiotemporal profiling of the microbiome at ecosystem-scale. These methods can identify spatial relationships among hundreds of microbes without requiring prior specification of target taxa. Additionally, these methods have shown strong correlations between spatial associations in stool and fecal samples, which enables efficient and cost-effective sampling for studies of changes in the spatial organization of the gut microbiome in the same individual over time. Protocols for these methods involve fixing samples in an acrylamide matrix, cryo-fracturing into particles, and passing particles through sizing filters, typically ∼40 microns in diameter. A bar-coding procedure is then performed, in which DNA amplicons from microbes in the same particle receive identical sequencing barcodes. Finally, amplicons are released from the particles and sequencing is performed on standard instruments, with bioinformatic processing producing a table of counts per particle for each taxa identified in the sample. Computational tools are essential for analysis of these data, given their high-throughput nature and particular features, including high-dimensionality, counts-based measurements, uneven amplification resulting in extremely variable numbers of reads per particle, and contamination from unencapsulated DNA mixing into particles^30^. However, existing tools have significant limitations. For instance, the most common approach in use binarizes the data, which discards potentially valuable quantitative information, and additionally restricts analyses to pairwise associations^11^. Clustering models have also been applied to these data^35^, which can detect multi-way interactions, but this approach does not address other important properties of these data, including its unique noise characteristics. Additionally, none of the existing methods model temporal changes or perturbations introduced to the ecosystems.

To enable characterization of microbiome spatiotemporal dynamics at ecosystem-scale, we developed a purpose-built computational tool, MCSPACE, for analyzing sequencing-based microbial co-localization data. MCSPACE, which we make available to the community as an open-source software package, introduces capabilities that address limitations of prior tools, namely: (1) characterization of spatially co-occurring groups of microbes, termed *assemblages*, to capture multi-way associations, (2) a custom noise model tailored for high-throughput spatial co-localization data, (3) automatic determination of changes in assemblage abundances over time and due to introduced perturbations. To evaluate our method, we generated the most comprehensive time-series of microbiome spatial co-localization to date, consisting of SAMPL-seq analyses of 21 fecal samples from a cohort of three mice subjected to a series of dietary perturbations conducted over a 76-day period. Using this dataset and an existing longitudinal SAMPL-seq dataset of the human gut microbiome, we demonstrate that MCSPACE outperforms state-of-the-art methods and uncovers biologically relevant spatiotemporal dynamics in both human and murine gut microbiomes.

## Results

### MCSPACE is an open-source computational tool for inferring spatiotemporal dynamics of microbiomes from high-throughput microbiome co-localization data

We implemented a custom, generative AI-based model for identifying, from sequencing-based co-localization data such as SAMPL-seq (**Figure 1A–C**), spatially coherent groups of microbes, termed *assemblages*, and detecting changes in the proportions of these assemblages over time and due to introduced perturbations. MCSPACE employs a Bayesian probabilistic technique that models the observed counts-based sequencing data as being generated by latent, or unobserved, random variables (**Figure 1D-F**). These variables are learned from data using an approximate inference method based on a generative AI technique, Variational Autoencoders^36,37^, which is scalable and flexible, enabling modeling tailored to the specific properties of the data. Further, this method provides estimates of confidence, allowing the user to robustly interpret inferences derived from noisy data.

**Figure 1.**
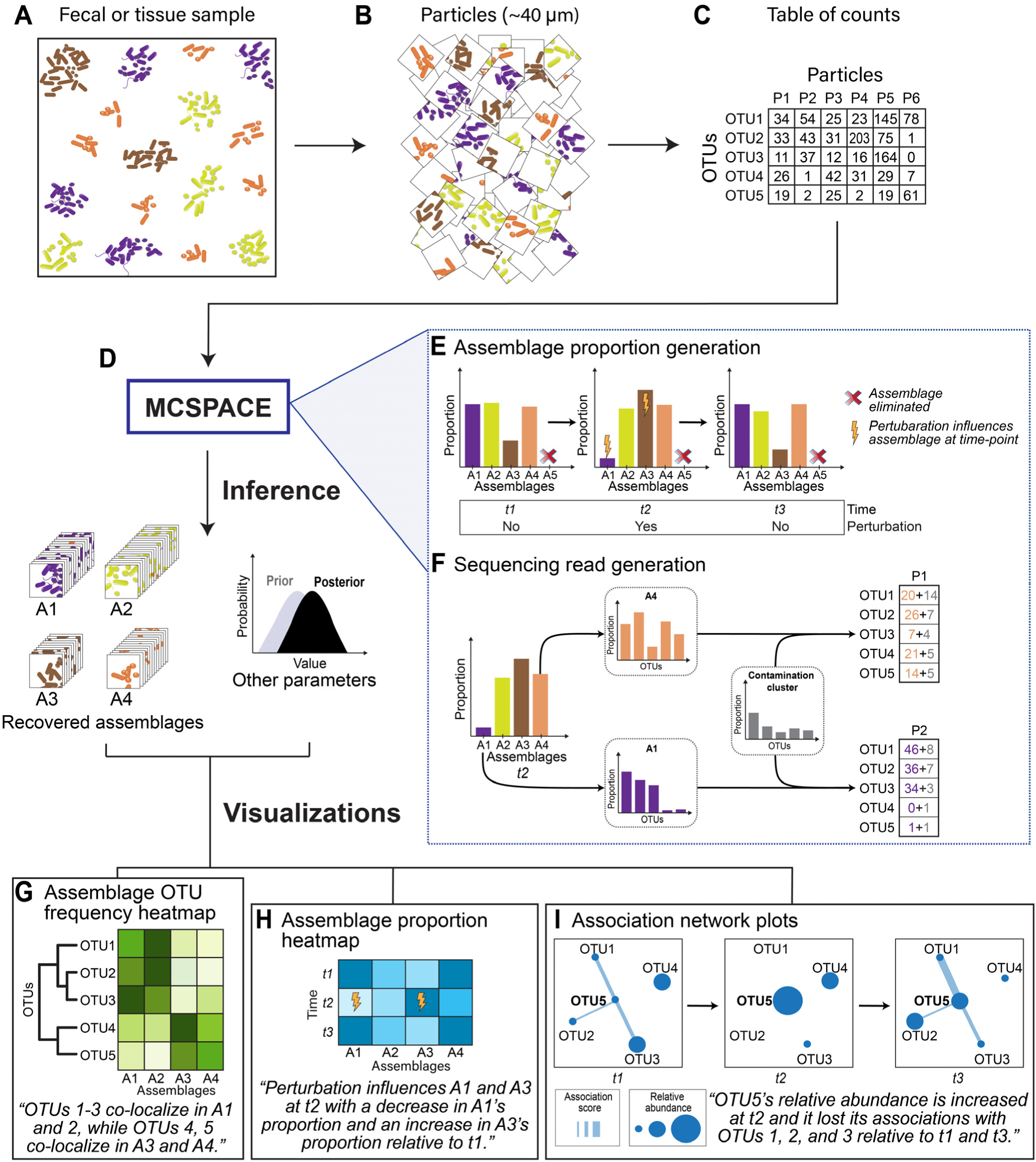
MCSPACE is a computational tool for inferring spatiotemporal dynamics of microbiomes from high-throughput sequencing-based co-localization data. This schematic depicts the flow from the physical input sample to the outputs produced by MCSPACE. **(A)** Fecal or tissue sample with spatial arrangements of microbes. **(B)** High-throughput co-localization data, such as SAMPL-seq, is generated by fragmenting the input sample into small (e.g., ∼40 um) particles. **(C)** DNA in particles is barcoded and then sequenced in bulk to produce a table with counts of Operational Taxonomic Units (OTUs) in each particle. **(D)** MCSPACE uses a purpose-built generative AI model to infer, from the input table of counts, microbial assemblages, or groups of recurrently spatially associated microbes, and changes in abundance of the assemblages over time and due to experimentally introduced perturbations. **(E)** and **(F)** depict the latent generative process assumed by MCSPACE. **(E)** Proportions of assemblages in each sample are modeled as potentially changing over time and with perturbations. Redundant assemblages or irrelevant changes with perturbations are automatically eliminated during inference. **(F)** Observed sequencing reads for a particle are generated by first picking an assemblage based on the assemblage proportions for the sample. Then, for each read in the particle, the model chooses to generate it either from the assemblage, or from a contamination cluster that accounts for unencapsulated DNA mixing into particles. Assemblages and the contamination cluster are modeled as frequencies of OTUs. MCSPACE outputs three types of visualizations of the inferred model: **(G)** assemblage OTU frequency heatmaps organized by phylogeny, **(H)** assemblage proportion heatmaps across time and depicting perturbation influences, and **(I)** association network plots that depict changes in abundance and associations for anchoring taxa of interest.

The MCSPACE model was purpose-built to capture key features of high-throughput microbiome co-localization data (**Figure 1E-F**). Observed counts of Operational Taxonomic Units (OTUs) in particles are assumed to arise from a set of latent assemblages that consist of frequencies of OTUs. MCSPACE automatically determines the number of assemblages using a Bayesian Variable Selection (BVS) technique, with a bias toward parsimony (e.g., the lowest number of assemblages warranted by the data). Proportions of assemblages are assumed to change over time (**Figure 1E**), and may also change with introduced perturbations, but with a bias toward no changes modeled again using a BVS technique. Observed reads for each particle are thus assumed to be generated by the following process (**Figure 1F**): (1) each particle probabilistically chooses the assemblage it comes from, according to the assemblage proportions for the sample, (2) each observed read chooses to either be generated from a *contamination cluster* (modeling mixing from unencapsulated DNA), or from the particle’s chosen assemblage, and (3) the read chooses which OTU it comes from, according to the OTU frequencies of either the assemblage or contamination cluster. See **Methods** for full details on the model and inference procedure.

Inputs to the open-source software package are: (1) a table of microbial sequencing read counts in each particle, in each sample, (2) a table of taxonomic labels for OTUs, and (3) metadata with a time-point index for each sample, and a flag for each time-point indicating whether a perturbation was applied. The software outputs a set of tables and visualizations to facilitate interpretation of its results by biologists, including visualizations of (**Figure 1G-I**): (1) members of assemblages, organized into phylogenetic trees; (2) the proportions of assemblages in each sample, and whether each assemblage has been identified as being influenced by each perturbation; (3) changes in spatial associations between taxa over time.

### Human and mouse SAMPL-seq spatial co-localization dataset compendium

For investigating spatiotemporal dynamics of the human gut microbiome, we used an existing longitudinal SAMPL-seq dataset of fecal samples collected from a healthy human donor over five consecutive days^31^ **(Figure 2A,B)**. Raw read counts obtained from the study were filtered for quality (see **Methods, Figure S1**). After quality filtering, the dataset consisted of 58 OTUs and 7055 particles (∼40 um in diameter), with a median of 1419 (interquartile range [IQR]=451) particles per sample **(Figures 2C, S2)**. **Figure 2D** visualizes the number of reads per particle (median = 843, IQR = 1828).

**Figure 2:**
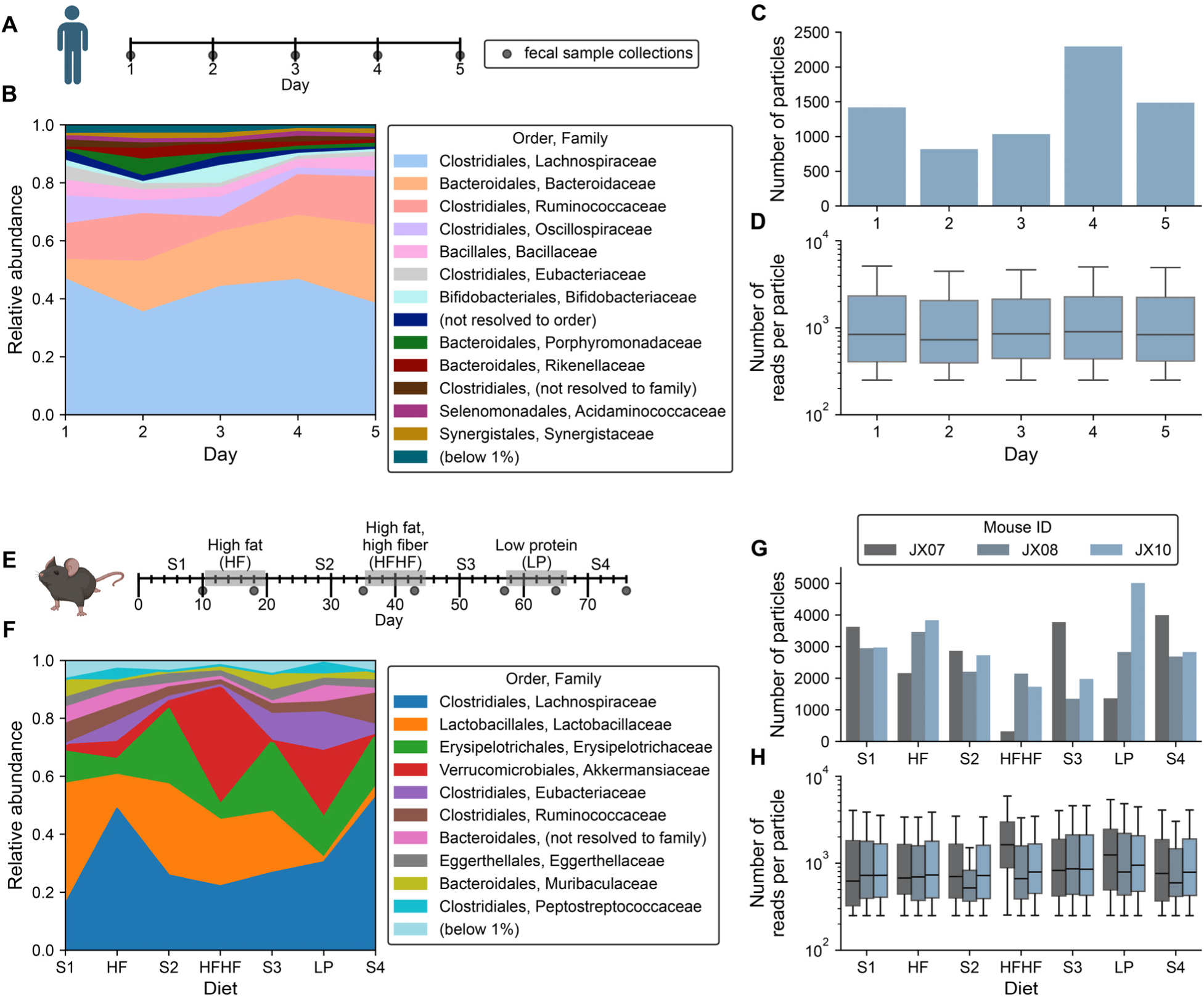
C**o**mpendium **of SAMPL-seq longitudinal datasets including new murine dataset with multiple dietary perturbations. (A-D)** Reanalysis of data from a study of gut microbiome spatial co-localizations in a healthy human participant (collected daily, *n*=5 fecal samples total). **(E-H)** Longitudinal murine gut microbiome spatial co-localization study (*n*=21 fecal samples total, collected from 3 mice). Dietary perturbations were selected to cause significant and differential changes to the composition of the microbiota. S1-4 = standard chow. **(B, F)** Microbial compositions of datasets at the Family level, aggregated over particles. For the mouse dataset, the mean proportions are shown over the animals. **(C, G)** Particle counts and **(D, H)** reads per particle distributions after quality filtering. Boxplot centers in **(D, H)** show medians, with boxes representing interquartile ranges (IQRs). Whiskers extend to data points within 1.5 × IQR.

To more comprehensively investigate microbiome spatiotemporal dynamics, we performed a study in mice **(Figures 2E, S3)**. A cohort of three C57BL/6 Jackson Laboratory (Jax) mice were subjected to defined dietary changes over a 76-day period. After mice were equilibrated on standard chow for 10 days, each dietary perturbation was introduced in sequence, with intervening periods of two weeks on standard chow for washout^7,18^ of the perturbation diet. We selected the perturbation diets–high-fat (HF), high-fat high-fiber (HFHF), and low-protein (LP)–for their known strong and distinct effects on microbial composition, with each differentially affecting microbes that preferentially utilize certain carbohydrate, fat, protein, and fiber sources^7,38,39^. Details on the composition of the diets are provided in **Table S1.** Fecal samples were collected from each mouse on each diet after equilibration, at days 10, 18, 35, 43, 57, 65, and 76 **(Figure 2E)**, yielding 21 samples for SAMPL-seq analyses **(Figure 2F)**. After quality filtering, the dataset **(Figures 2G, S3)** consisted of 74 OTUs and a median of 2829 particles (∼40 um in diameter) per sample (IQR=1319). **Figure 2H** visualizes the number of reads per particle (median = 745; IQR = 1390).

### MCSPACE accurately recovers spatial associations from semi-synthetic data

MSPACE’s utility hinges on its ability to accurately recover spatial associations from high-throughput data. Because no comprehensive source of ground-truth microbiome associations exists to assess this capability, we employed a bootstrapping-type procedure that generates semi-synthetic data from the MCSPACE model, while preserving distributional properties of real SAMPL-seq data. Briefly, data were simulated using the 34 latent assemblages inferred on the first time-point of the human dataset. The human dataset was chosen as it contained more OTUs per particle (**Figure S2B**), and thus presents a more challenging task for recovering spatial associations. We simulated datasets that varied the number of particles or reads per particle, to assess how these features of SAMPL-seq data could affect MCSPACE’s performance. Values for the numbers of particles and reads per particle were chosen to mimic variability of dataset sizes seen in SAMPL-seq runs, with values of 250, 500, 1000, 2500, and 5000 used, spanning approximately an order of magnitude below and above the values observed in real data. Each feature was varied individually, while the other was kept constant (set to the value inferred on real data). See **Methods** for details on our data simulation procedure.

We additionally used the semi-synthetic data to evaluate computational methods that have previously been used for analyzing high-throughput microbial spatial co-localization data. The semi-synthetic data, which were simulated from the MCSPACE model itself, obviously cannot be used to draw conclusions about the absolute performances of other methods on real data. However, these analyses can provide insights into how features of the data that vary with experiments, such as the depth of sequencing, could affect results for different computational methods in a relative sense. Two of the methods^30,31^ detect pair-wise associations by binarizing data then using either Fisher’s exact test or generating a null distribution based on the SIM9 algorithm from the ecology literature^40^, to determine statistical significance. A third method^35^, Gaussian Mixture Models (GMMs), a standard clustering approach, identifies clusters of particles and is thus capable of finding multi-way interactions. Note that a purpose-built GMM model including directional gradients over clusters^35^ has also previously been applied to high-throughput microbiome spatial co-localization data; we evaluated this method on real data, but were unable to do so on the semi-synthetic data because the method failed to converge on more than ten clusters, making fair comparison to the other methods infeasible.

#### Detecting pairs of spatially co-associated microbes

Pair-wise associations are a frequent unit of analysis in microbial ecosystems^18,41–43^, so we wanted to assess whether the existing algorithms^30,31^ for recovering these types of associations from high-throughput microbiome spatial co-localization data could do so in the setting of underlying multi-way spatial associations, which occur in real microbial communities^11^. We evaluated performance using the area under the receiver operator curve (AUC), for the ability to detect pairs of associations, and found that both the Fisher’s exact and SIM9 methods did fairly well, achieving AUCs of around 0.80 and 0.75, respectively (**Figure 3A,B**). MCSPACE did significantly better (Wilcoxon rank sum test, *p*<0.001), with median AUC differences ranging from 0.17 to 0.20 when compared to Fisher’s exact test, and 0.23 to 0.24 when compared to the SIM9 algorithm. None of the methods showed substantial improvements with increasing numbers of particles or read depths, although positive trends were most clearly evident for the numbers of particles. These results demonstrate that pairwise spatial associations can be detected from data with underlying multi-way associations, and suggest that although prior methods perform reasonably well on this task, there is room for improvement.

**Figure 3:**
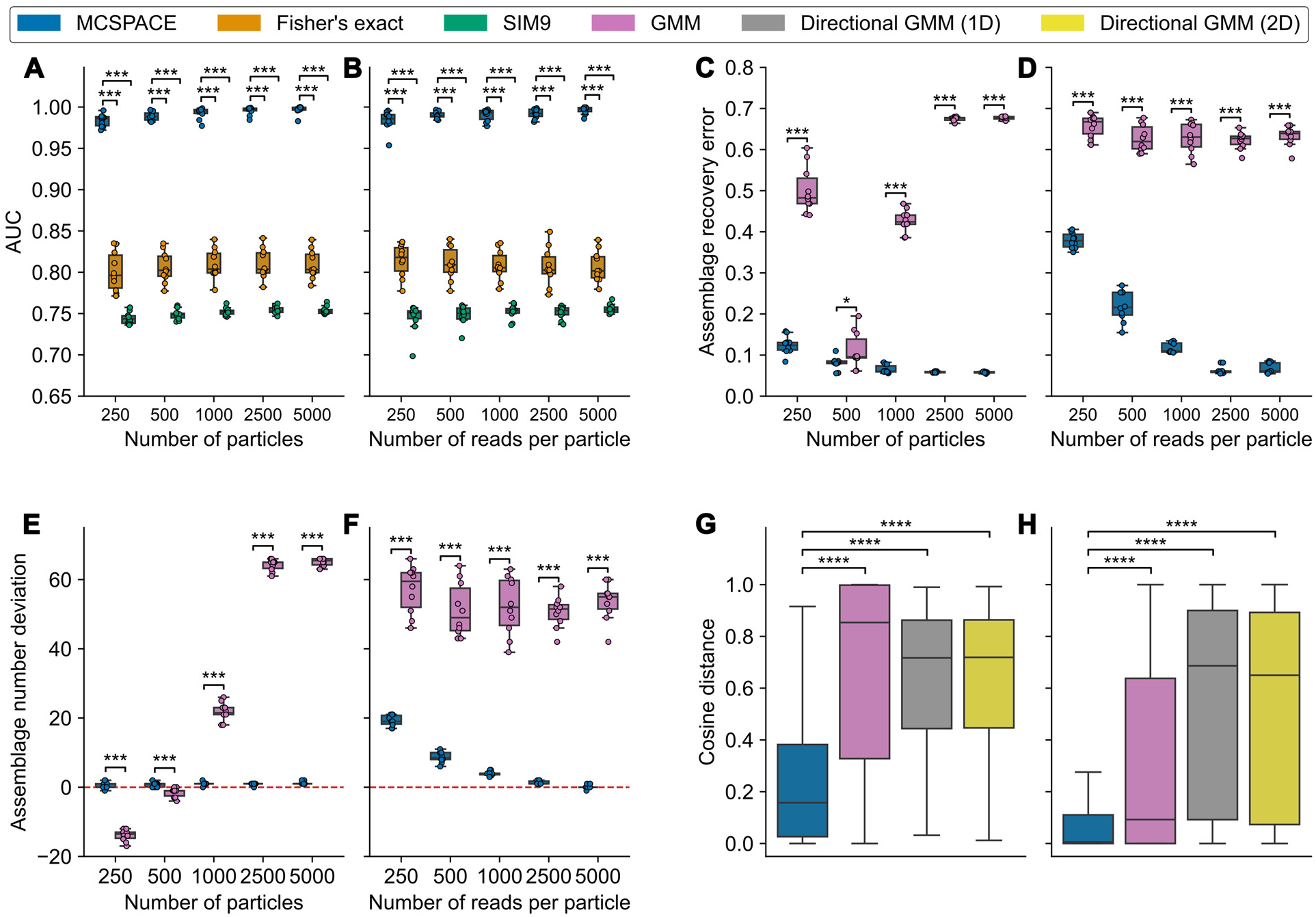
M**C**SPACE **accurately recovered spatial associations in semi-synthetic data and significantly outperformed comparator methods in predicting held-out real data.** MCSPACE and five existing methods were evaluated: Fisher’s exact and SIM9, which detect pairwise interactions, and three types of Gaussian Mixture Models (GMMs), which infer multiway assemblages. **(A-F)** Semi-synthetic data was simulated from the MCSPACE model inferred on real data. Methods were assessed for their ability to recover ground-truth information in three tasks: **(A,B)** Detecting co-associated pairs of microbes, assessed with area under the receiver operator curve (AUC) (higher values indicate superior performance), **(C,D)** Recovering the correct frequencies of OTUs in assemblages, assessed using assemblage recovery error (lower values indicate superior performance), and **(E,F)** Inferring the correct number of assemblages, assessed by subtracting the inferred value from the correct value (values nearer to zero indicate superior performance). Methods were benchmarked on real human **(G)** and mouse **(H)** data for their ability to predict held-out sequencing reads using 5-fold cross-validated cosine distance as the evaluation metric (lower values indicate superior performance). Boxplots show results from 10 simulated replicates for synthetic data or all particles post-filtering in real data, (*n*=7055 for human and *n*=56,848 for mouse). Central lines indicate medians with boxes representing interquartile ranges (IQRs). Whiskers extend to data points within 1.5 × IQR. Statistical significance was assessed with a Wilcoxon rank sum test followed by Benjamini-Hochberg correction for multiple hypothesis testing (*, p<0.05; **, p<0.01, ***: p<0.001, ****: *p*<0.0001).

#### Recovering spatial assemblages

We next assessed whether models (GMMs and MCSPACE) capable of recovering multi-way spatial associations, or assemblages, could do so accurately. To assess this capability, we developed an assemblage recovery error metric, which matches inferred and true assemblages and calculates distances between matched assemblages (**Figure S4**); see **Methods** for details. GMMs performed poorly overall on this task, showing no clear improvement with more particles and only slight improvement with increasing read depth (**Figure 3C,D**). In contrast, MCSPACE performed significantly better (Wilcoxon rank sum test*, p*<0.001), up to 9-fold, and showed steady gains with increasing numbers of particles and read depths. To gain further insight into these differences in performance, we evaluated how well GMMs and MCSPACE could recover the underlying number of assemblages (**Figure 3E,F**). GMMs generally substantially overestimated the number of assemblages, although the method underestimated the number of assemblages in the setting of low numbers of particles; there was no clear trend of improving performance with additional data. In contrast, MCSPACE’s performance was significantly better (Wilcoxon rank sum test*, p*<0.001), with performance steadily improving with more data, especially with increasing numbers of reads per particle. These results demonstrate that GMMs have difficulty recovering microbiome spatial assemblages from data with realistic parameters, and there thus may be advantages to a purpose-built model such as MCSPACE.

### MCSPACE outperformed comparator methods on real data from both humans and mice

To assess MCSPACE’s and comparator techniques’ performance on real data, we evaluated the ability of the methods to predict held-out data. This is a common benchmark for generative models in the machine learning literature (e.g., document completion tasks in topic models^44^), which assesses models’ abilities to capture distributional properties of real data accurately. We assessed performance via a five-fold cross-validation procedure, with each fold trained on 80% of particles containing all their sequencing reads and 20% of particles containing 50% of their reads (with 50% held-out), with the task of predicting the held-out reads. This procedure, of not holding out the entirety of individual units of analysis (particles, in our case), is standard in the machine learning literature and is necessary to provide appropriate context for the model to reconstruct the held-out data. We used the cosine distance between predicted and held-out data as the performance metric (lower values indicated superior performance). See **Methods** for details on the evaluation procedure.

For this task, we compared MCSPACE against standard and directional GMMs^35^, which are also generative models. For each cross-validated fold, MCSPACE was run with a capacity of *K* = 100 assemblages. The standard GMM was run for a range of 2 to 100 assemblages, and the best model was selected post hoc as described in **Methods**. Directional GMMs failed to converge when the number of assemblages exceeded 10 and were therefore only run up to 10 assemblages. Both standard and directional GMMs inherently assume data is Normally distributed and thus model transformed data, rather than the actual data, which consists of sequencing counts. Also, these methods use a post hoc procedure to control model sparsity, unlike MCSPACE, which controls sparsity directly as part of its Bayesian model.

MCSPACE significantly outperformed all comparator models on this task on our compendium containing both human and mouse datasets (Wilcoxon rank sum test, *p*<0.0001, **Figure 3G,H**). On human data, MCSPACE had a median cosine distance of 0.16, whereas the GMM models ranged from 0.71 to 0.85; on mouse data, MCSPACE’s median cosine distance was 0.005 compared to GMM models ranging from 0.09 to 0.69. These results demonstrate that MCSPACE, which was purpose-built to capture key aspects of real data such as modeling sequencing counts and particle contamination, outperforms models without these capabilities in accurately reproducing real high-throughput microbiome co-localization data.

### Spatiotemporal dynamics of human and murine gut microbiomes

#### MCSPACE elucidated both dynamic and persistent associations in the human gut microbiome

To gain insights into spatiotemporal dynamics of the human gut microbiome, we applied MCSPACE to the dataset from our compendium profiling fecal samples from a single healthy individual over five consecutive days. MCSPACE identified 58 OTUs assorting into 40 spatial assemblages, with assemblages dominated by abundant bacteria such as *Phocaeicola dorei* OTU3, present in 22 of 40 (55%) assemblages at ≥5% abundance, and *Agathobacter rectalis* OTU1, present in 14 of 40 (35%) assemblages at ≥5% abundance (**Figure S5**). These findings are consistent with previous research^31^ that identified these taxa as central “spatial hubs” in the gut. The widespread presence of *Phocaeicola dorei* in assemblages is also consistent with the role of *Bacteroidaceae* as generalists capable of occupying diverse spatial niches^45^.

To assess the dynamic versus persistent nature of assemblages, we examined their temporal proportions and identified the assemblages exhibiting the most and least variability. We defined an assemblage as *variable* if it was at least 3-fold more abundant at one-time point compared to all other time points, or *persistent* if the standard deviation of its log proportions over time was less than 1%. Using this definition, we identified three variable and four persistent assemblages (**Figure 4**). These assemblages exhibited distinct sets of abundant OTUs, with no OTUs overlapping between them.

**Figure 4:**
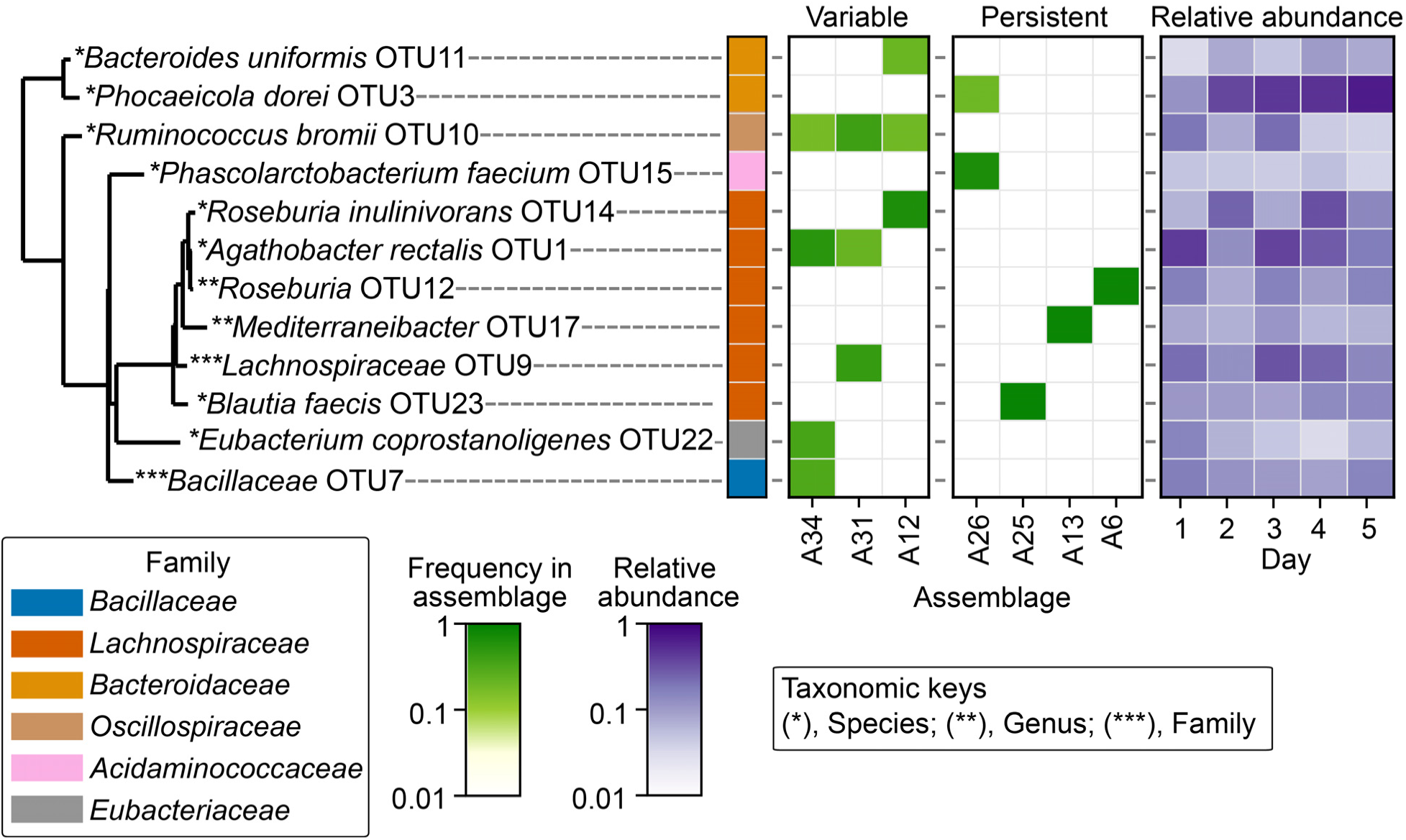
M**C**SPACE **analysis of longitudinal high-throughput co-localization data of the human gut microbiome revealed both dynamic and persistent spatial associations.** Assemblages with ≥3-fold abundance increase at one-time point compared to all other time points were classified as variable; assemblages with the variance of their log proportions over time <1% were classified as persistent. OTUs that are present in either variable or persistent assemblages with >5% frequencies are shown. Variable assemblages primarily contained taxa capable of degrading resistant starches (e.g., *Agathobacter rectalis* and *Ruminococcus bromii*) and/or plant fiber polysaccharides (e.g., *Roseburia inulinivorans* and *Bacteroides uniformis*). In contrast, persistent assemblages contained mostly taxa that use nutrients either from cross-feeding interactions (e.g., *Phascolarctobacterium faecium* is a succinate-consumer and *Phocaeicola dorei* is a producer) or from ubiquitous diet-or host-derived sources (e.g., *Blautia*, *Mediterranibacter* and *Roseburia* species produce butyrate from a variety of substrates that are common in the Western diet, as well as from host-derived mucin).

The variable assemblages primarily contained taxa capable of degrading resistant starches and/or plant fiber polysaccharides, with many of these taxa previously having been shown to be responsive to meal-to-meal changes in dietary fiber^46–50^. Assemblages A34 and A31 both peaked on day 1, with A34 exhibiting the largest fold increase, over 4.5-fold relative to other days. Both assemblages contained *Agathobacter rectalis* OTU1 and *Ruminococcus bromii* OTU10, which are known resistant starch degraders^48^. Notably, *A. rectalis* has been shown to have enhanced growth on resistant starch when co-cultured with *R. bromi* in vitro^51^. This raises the possibility that the assemblages with *A. rectalis* and *R. bromi* could reflect co-colonization of diet-derived starch granules^52,53^ and syntrophic interactions^51^, with variability of these assemblages possibly due to fluctuations of resistant starch intake by the host. Assemblage A12, which peaked on day 2, also contained *R. bromii* OTU10, along with *Roseburia inulinivorans* OTU14 and *Bacteroides uniformis* OTU11 (both degraders of plant fiber polysaccharides^49,50^). *Bacteroides spp.* are known nutritional generalists^45^, with the well-studied type strain *Bacteroides thetaitaomicron* having been shown to attach to food particles through glycan-specific outer-membrane binding proteins during carbohydrate scavenging behavior^54^. Additionally, motility mechanisms involving inducible flagellar genes that enhance its access to carbohydrates^49^ have previously been reported in *R. inulinivorans*, providing a potential explanation for its ability to readily reorganize into spatial assemblages with dietary shifts. Overall, our findings demonstrate that assemblages of spatially co-localized bacteria in the human gut with highly variable abundances are dominated by organisms that utilize complex carbohydrates that are not as common in Western diets as simpler nutrients, suggesting the hypothesis that the variable levels of these assemblages may be driven by transient consumption of resistant starches and plant fiber by the host.

Persistent assemblages, in contrast, contained mostly taxa that use more broadly available nutrients. The least variable assemblage, A26, was dominated by *Phascolarctobacterium faecium* OTU15 (55%) and *Phocaeicola dorei* OTU3 (8%). *P. faecium* is a known succinate consumer^55,56^, while *P. dorei* OTU3 produces succinate during central metabolism^57^, suggesting a cross-feeding interaction between them. The other three persistent assemblages, A25, A13, and A6, were each dominated by a single taxon from the *Lachnospiraceae* family: *Blautia faecis* in OTU23, *Mediterranibacter* in OTU17, and *Roseburia* in OTU12. These taxa are abundant in the gut and produce butyrate from a variety of substrates that are common in the Western diet, as well as from host-derived sources including mucin^58–62^. Our findings thus demonstrate that spatially co-localizing assemblages of bacteria in the human gut with nearly constant abundances are dominated by organisms that can use nutrients that may be provided by various stable sources including other microbes, common dietary components, or the host.

#### MCSPACE uncovered diet-induced spatial redistributions in the murine gut microbiome

To obtain greater insights into the impact of dietary nutrients on the spatiotemporal dynamics of the gut microbiome, which was suggested by the human data but could not be further investigated because that study used an *ad libitum* diet, we applied MCSPACE to the mouse SAMPL-seq dataset with controlled dietary perturbations. This analysis identified 74 OTUs assorted into 63 spatial assemblages (**Figure S6**). Of these assemblages, MCSPACE identified strong evidence^63^ (Bayes Factor > 10) that the abundances of 14 were perturbed by at least one of the three dietary interventions compared to their abundances on the preceding standard chow diet period (**Figure S7**): five assemblages were perturbed by the HF diet, three by the HFHF diet, and eight by the LP diet, with two assemblages perturbed by multiple diets (A18 by both the HF and LP diets; A3 by both the HFHF and LP diets). These findings indicate that dietary changes can have significant effects on the spatial organization of the gut microbiome, with both specific and broad effects evident.

Given the observed overall strong effects of diet on spatiotemporal dynamics of the gut microbiome, we next sought to gain a more detailed understanding of how spatial associations among microbes reorganize with diet. To conduct this analysis, we developed a pairwise metric that scores the degree of spatial co-localization on each diet between an anchoring taxon and those it associates with (see **Methods**) and maps that visualize gains and losses of associations across the diets (**Figure 5**). Anchoring taxa were selected as those with a minimum relative abundance of 5% on at least three dietary intervals, which yielded two taxa: *Akkermansia* OTU2 and *Lactobacillus* OTU1.

**Figure 5:**
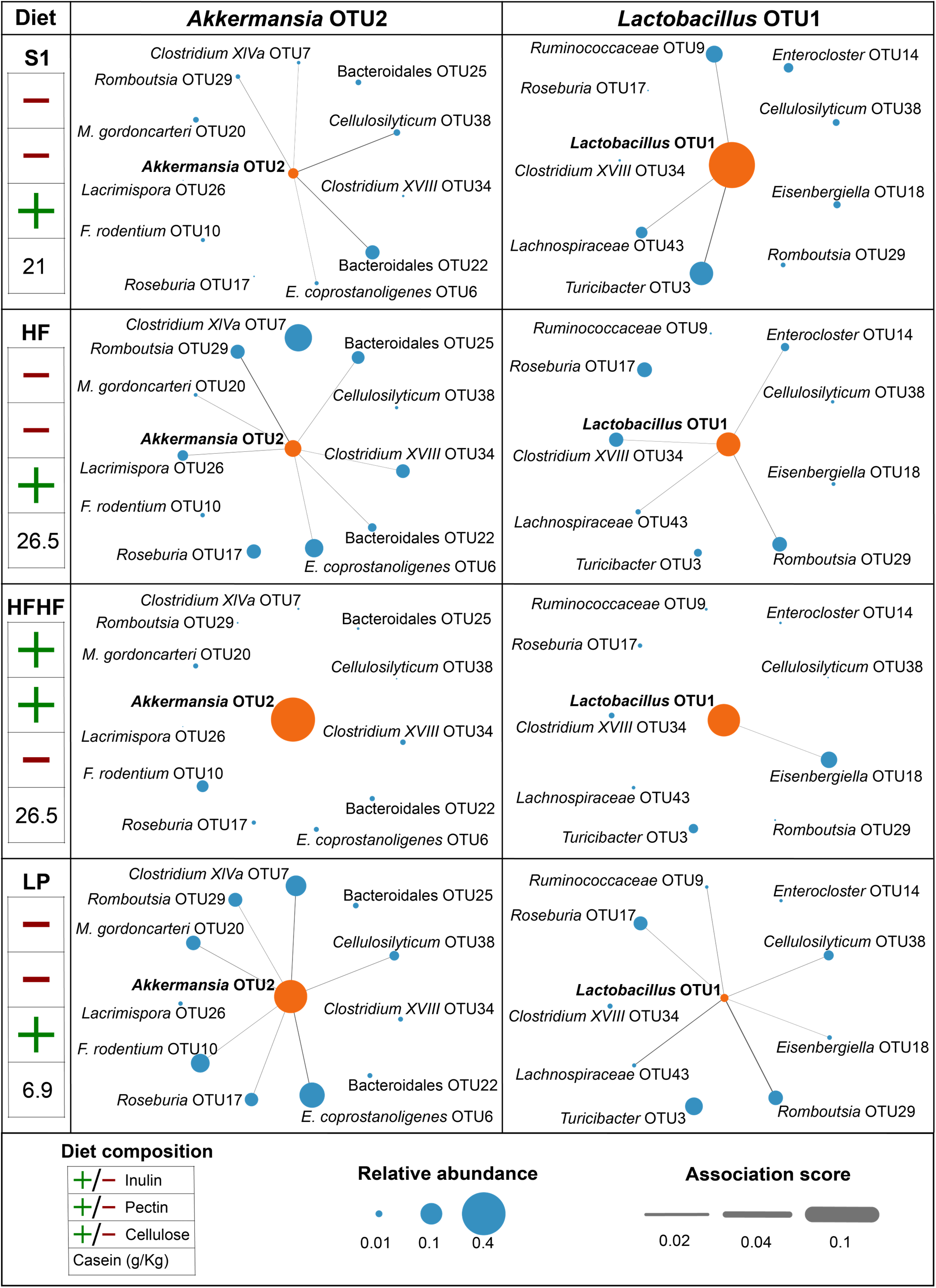
M**C**SPACE **analysis revealed nutrient-induced changes in spatial associations in the murine gut microbiome in our study with controlled dietary perturbations.** Taxa to anchor differential spatial association mapping analyzes were selected as those with a ≥5% relative abundance on ≥3 dietary intervals, yielding two taxa: *Akkermansia* OTU2 and *Lactobacillus* OTU1. Maps depict associations on S1 (standard chow), HF (high fat), HFHF (high fat, high fiber), and LP (low protein) diets. Node sizes indicate relative abundances of taxa and edge widths indicate the strengths of spatial associations (<0.01 not shown). *Akkermansia* OTU2’s spatial associations shifted significantly with the fiber type in the diet, persisting on insoluble cellulose (S1, HF, LP) and disappearing on the soluble fibers inulin and pectin (HFHF). *Lactobacillus* OTU1 similarly lost associations with soluble fibers in the diet (HFHF). Additionally, when *Lactobacillus* OTU1’s preferred fermentation substrate casein was significantly reduced (LP), the taxon decreased in abundance while increasing its spatial associations, suggesting that *Lactobacillus* OTU1 may engage in cross-feeding with taxa such as *Cellulosilyticum* OTU38 (a degrader of dietary cellulose) when preferred nutrients are not readily available.

The first anchoring taxon, *Akkermansia* OTU2, showed extensive reorganization of its spatial associations (**Figure 5**) with diet. Although this taxon had only modestly higher relative abundance on the HF diet (5.8%, compared to 2.2% on standard chow), many other taxa had spatial associations with it than they did on standard chow, including Bacteroidales OTU25, *Muribaculum gordoncarteri* OTU20, *Clostridium XVIII* OTU34, *Lacrimispora* OTU26, and *Romboutsia* OTU29. Interestingly, on the HFHF diet, which uses inulin and pectin as fiber sources versus the other diets that use cellulose, *Akkermansia* OTU2’s relative abundance was dramatically higher (40%), but other taxa lost all spatial associations with it. In contrast, although *Akkermansia* OTU2’s relative abundance was also high (23%) on the LP diet, which has reduced protein content and increased sucrose content relative to the other diets, multiple taxa retained their associations with *Akkermansia* OTU2. Soluble fibers, such as inulin and pectin, have been shown to be readily fermented to produce short-chain fatty acids by a broad range of microbes^64–66^, including *Akkermansia*. In contrast, insoluble fibers such as cellulose are not readily metabolized by most microbes, but can stimulate production of host glycans such as mucin^67,68^, which indirectly benefits *Akkermansia*, a mucin degrader^69^. Our findings thus suggest the hypothesis that insoluble fibers available from the diet may promote cross-feeding and gains in microbial spatial co-localizations organized around *Akkermansia* as a hub organism that can degrade host glycans and produce a variety of metabolic byproducts useful to other bacteria^70–72^, whereas insoluble fibers in the diet can promote independent growth^64–66^ of bacteria and loss of spatial co-localizations with *Akkermansia* when the reliance on cross-feeding is lessened.

The second anchoring taxon, *Lactobacillus* OTU1, also showed extensive reorganization of its spatial associations (**Figure 5**) with diet. This taxon had associations with three taxa on standard chow and four on the HF diet. Similar to *Akkermansia* OTU2, *Lactobacillus* OTU1 lost its associations on the HFHF diet. Interestingly, *Lactobacillus* OTU1’s relative abundance was substantially lower on the LP diet (1.3%, versus 41% on standard chow, 11% on the HF diet, and 20% on the HFHF diet), while it exhibited the most spatial associations on this diet, including with members of the *Lachnospiraceae* family (*Lachnospiraceae* OTU43, *Roseburia* OTU17, *Cellulosilyticum* OTU38, *Eisenbergiella* OTU18), as well as with *Romboutsia* OTU29 and *Ruminococcaceae* OTU9. *Lactobacillus* species have a preference for fermenting casein compared to other gut bacteria^73^, and although all the diets in our experiments used casein as a protein source, the LP diet contained approximately one third the casein of the other diets. When casein is not available, *Lactobacillus* species can also efficiently utilize other energy sources, including soluble fibers such as inulin and pectin^66,74^. Our findings thus suggest the hypothesis that *Lactobacillus* OTU1 grows relatively independently in the gut when casein and soluble fibers are readily available in the diet (i.e., on the HFHF diet), whereas it co-localizes to engage in cross-feeding interactions when preferred energy sources are less available, such as interacting with *Cellulosilyticum* OTU38, a degrader of dietary cellulose^75^, on the LP diet.

## Discussion

We have introduced MCSPACE, an open-source software package that implements a custom generative AI-based method for extracting interpretable insights from high-throughput spatiotemporal microbiome data. To evaluate our method, we compiled a compendium of human and mouse spatiotemporal data, including a new murine dataset that we generated, which is the most comprehensive microbiome co-localization dataset to date. Using semi-synthetic data, we demonstrated that MCSPACE could accurately recover spatial associations among taxa, and on the compendium of real datasets, we showed that MCSPACE significantly outperformed state-of-the-art methods in predicting held-out data. Moreover, we showed that MCSPACE discovered interpretable spatial microbial assemblages, including temporally transient and persistent assemblages in the human gut microbiome and “hub” taxa that exhibit shifting spatial associations in the mouse gut microbiome in response to dietary changes.

Our computational model was purpose-built for its tasks, and thus offers distinct advantages over generic methods. Specifically, MCSPACE is a generative Bayesian method, employing a principled probabilistic framework that directly models irregular, sparse, and noisy counts-based data. In contrast, state-of-the-art comparator methods do not directly model the underlying data, but instead transform it using techniques such as binarization and log-ratio functions, which introduce distortions of the data and lose important information about its variability^76^. Our Bayesian method also has the advantage of producing quantitative measures of the uncertainty of its estimates, including the numbers of assemblages and the influences of perturbations, which allows the user to make informed decisions about which relationships warrant further investigation versus those that are more likely to be spurious. Indeed, the advantages of our approach were borne out by our superior benchmarking results. A challenge with Bayesian methods, such as ours, is that despite their strong capabilities, they are computationally costly to infer. We addressed this challenge by developing an efficient inference algorithm that approximates the MCSPACE model using generative AI techniques, while maintaining the highly interpretable structure of our model.

Application of MCSPACE to our human and mouse dataset compendium demonstrated our method’s ability to extract biologically meaningful results from these complex and noisy data. The assemblages and perturbation effects inferred by MCSPACE suggest that both temporally dynamic and persistent communities of microbes occur in the gut, and that the transience or durability of assemblages can depend on not only the particular taxa involved, but also the environmental context of the ecosystem (e.g., host diet). Variability in the diet may especially affect bacteria with strong nutrient preferences and that localize on food particles in the lumen^52–54^. In contrast, bacteria engaged in enduring cross-feeding interactions^55–57^ or that can assimilate common resources and grow in the gut environment with relative metabolic independence^58^ may form the most stable assemblages. We gained further insights into these phenomena through our mouse experiments with controlled dietary perturbations, including our finding that *Akkermansia* and *Lactobacillus*, taxa with marked nutrient preferences, dramatically changed their spatial associations with shifts in carbon and nitrogen sources. Overall, our results demonstrated MCSPACE’s power to characterize complex and dynamic spatial co-localization within the gut microbiome and to suggest specific hypotheses about the influence of host diet on ecological relationships among microbes.

Our study had several limitations, which suggest multiple directions for future work. First, the SAMPL-seq method only generates amplicons covering 69 bases of the 16S rRNA gene, which limits taxonomic resolution. Improvements in the method could yield longer reads and possibly profile additional genes, which would increase ability to differentiate species or even strain-level variations in spatial co-localization. Second, each dataset in our compendium was generated to characterize a single range of particle sizes. MCSPACE could be readily extended to handle multiple particle sizes, and with such data available, we expect our method would increase its overall accuracy in recovering spatial assemblages as well as provide new insights into assemblages at larger scales. Third, our mouse experiments employed a relatively limited set of perturbations. We chose these dietary perturbations because they caused large shifts in the microbiome, and we indeed saw dramatic changes in spatial co-localizations that could be attributed to microbial preferences for carbon and nitrogen sources. In future work, it would be interesting to explore a wider variety of diets and to quantitatively vary dietary components to establish the relationship between specific nutrients and microbial spatial structuring. Additionally, it would be interesting to study the effects of a broader range of perturbations, such as antibiotics, phages, and host pathogens. Fourth and finally, although the data we trained MCSPACE on, SAMPL-seq, has the advantage of being high-throughput, it has the disadvantage of lacking in situ spatial context. In future work, it would be interesting to incorporate in situ data, such as MERFISH^77,78^, GeoMX^79^, or Xenium^80^, as additional modalities informing assemblage membership. Data capable of spatially resolving metabolites or cell surface molecules, such as MALDI MSI^81^, would be particularly interesting to incorporate. These data could provide direct support for hypotheses generated by MCSPACE, including cross-feeding relationships that exist in the complex gut milieu but may be difficult to replicate in vitro.

In conclusion, our work provides new resources for characterizing spatiotemporal dynamics of the microbiome at ecosystems-level scale. We introduced the MCSPACE software, which makes sophisticated probabilistic AI capabilities available to researchers studying these complex ecosystems. Further, we have generated a comprehensive spatiotemporal dataset of the mouse gut microbiome that includes dietary perturbations, which we believe will serve as a valuable benchmarking and analytical resource for the community. Overall, our work provides a foundation for analyzing the spatial organization of microbiomes, which holds promise for unraveling the mechanisms of microbe-microbe and host-microbe interactions and developing means to harness the microbiome to promote health and ameliorate disease in the host.

## Acknowledgments

This work was supported by NSF MTM2 2025512, NIH NIGMS R01GM130777, NIH NIGMS R35GM149270, The Massachusetts Life Sciences Center, and the Brigham and Women’s Hospital President’s Scholar Award. G.U. was supported by the HHMI Hanna H. Gray Postdoctoral Fellowship (GT15182).

## Author contributions

Conceptualization, G.K.G. and H.H.W.; Methodology G.K.G. and G.S.U.; Software G.S.U; Validation G.S.U.; Formal Analysis G.S.U.; Investigation G.U., M.R., T.M. and J.L.; Resources G.K.G and H.H.W; Writing – Original Draft G.S.U, I.Y.K. and G.K.G; Writing – Review & Editing G.S.U, G.U., M.R. I.Y.C., T.M., J.L., H.H.W. and G.K.G.; Visualization G.S.U. and I.Y.K.; Supervision G.K.G. and H.H.W; Funding Acquisition G.K.G. and H.H.W.

## Methods

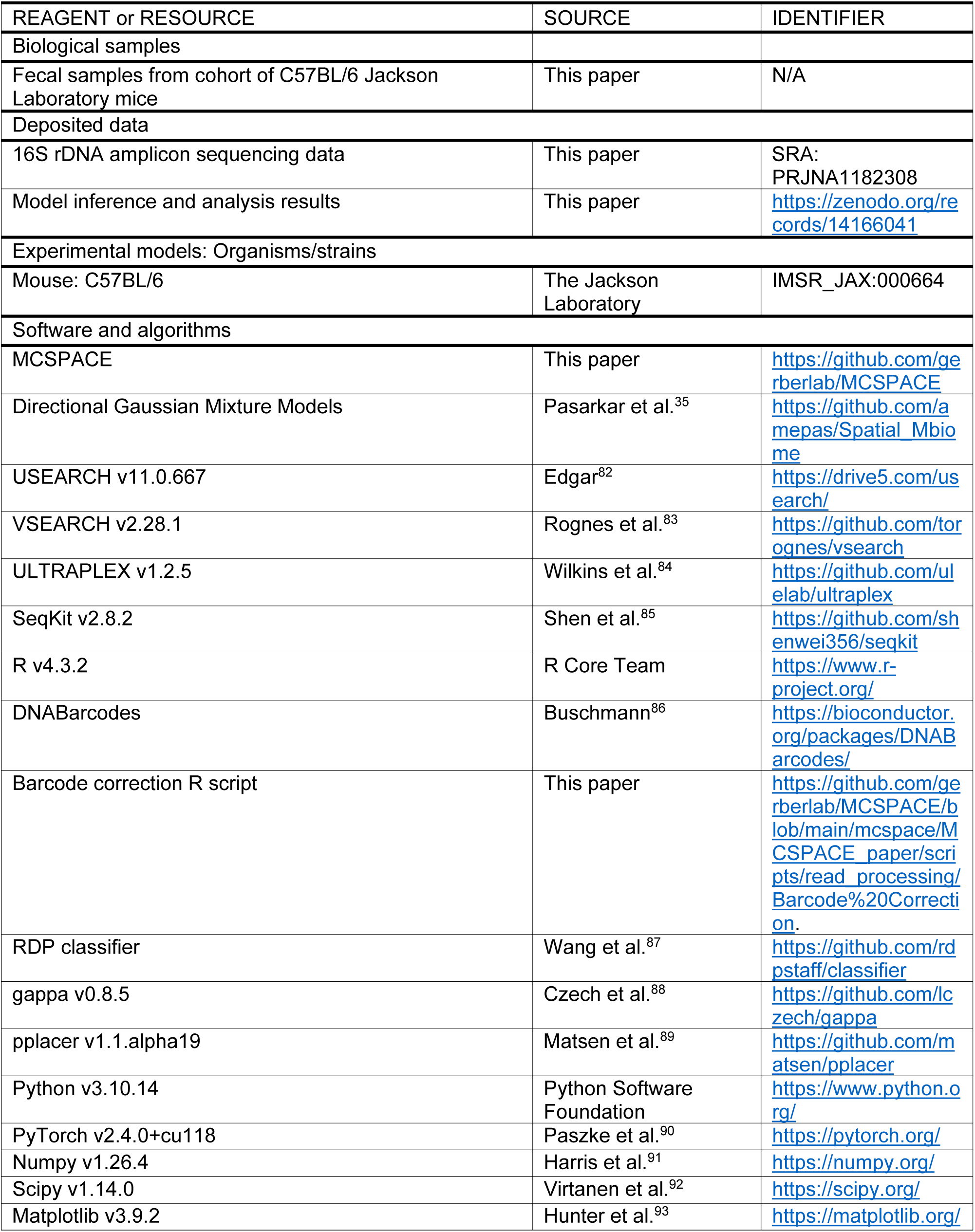
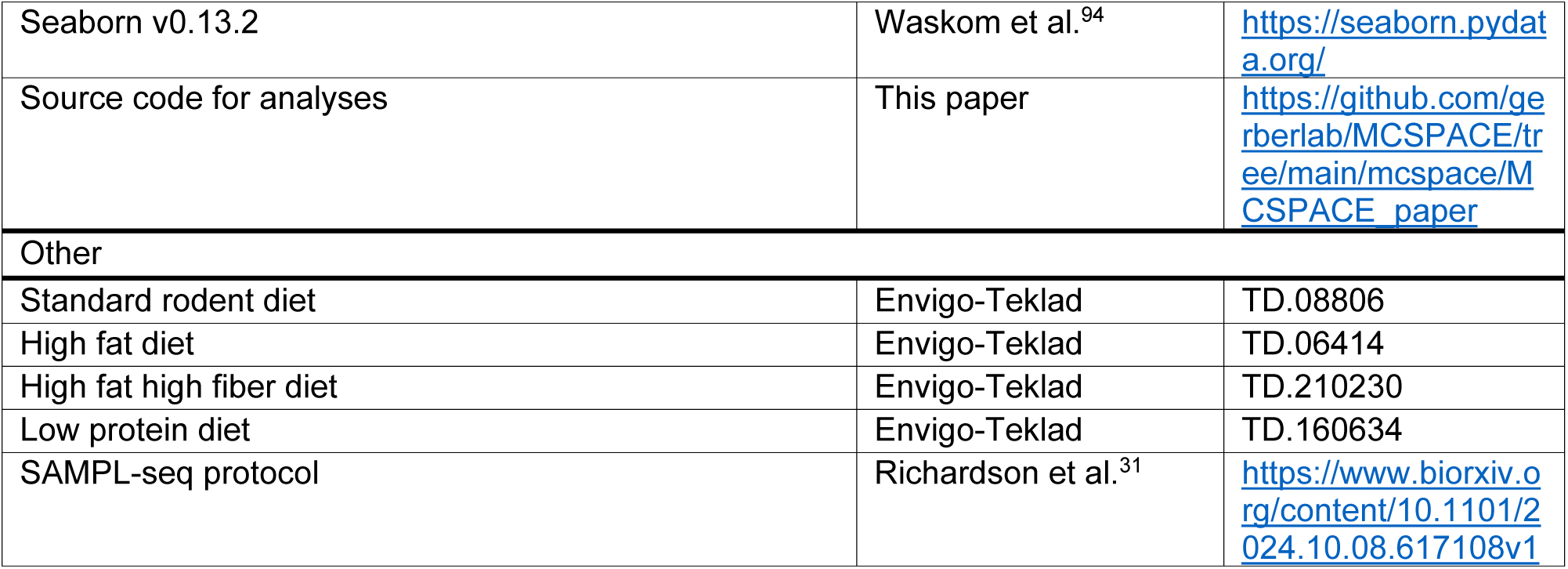
KEY RESOURCES TABLE.

## RESOURCE AVAILABILITY

### Lead contact

Further information and requests for resources and reagents should be directed to and will be fulfilled by the lead contact, Georg K. Gerber.

### Materials availability

This study did not generate new unique reagents.

### Data and code availability

- Raw sequencing data is available through SRA, and accession numbers are listed in the key resources table.
- The MCSPACE tool is publicly available with an open-source license on GitHub: https://github.com/gerberlab/MCSPACE. Code and processed data needed to generate figures from this study, Python notebooks with instructions for data processing, and all data necessary to run MCSPACE on example datasets are included. Additional files containing inference results from running the models are available on Zenodo repositories. Links to repositories are listed in the key resources table.
- All information required to reanalyze the data is available on GitHub and through the Zenodo link.

## EXPERIMENTAL MODEL AND STUDY PARTCIPANT DETAILS

### Human study participants

No human subjects research was conducted as part of this study.

### Mouse experiments

Experiments were conducted under Columbia University Medical Center Institutional Animal Care and Use Committee protocol AC-AABD4551. A cohort of three 6–8-week-old female C57BL/6 mice purchased from Jackson Laboratory were used in the experiments. The mice were co-housed and equilibrated for a period of 10 days on standard chow (Teklad custom diet, TD.08806) before undergoing a series of three dietary perturbations: high fat (HF), high fat high fiber (HFHF), and low protein (LP) diets, in that order. The HF and HFHF perturbations lasted for 10 days, and the LP perturbation lasted for 9 days, each followed by two-week periods off perturbations, on standard chow. For the HF perturbation, Teklad custom diet TD.06414 (60 kcal% of fat) was used. For the HFHF diet, Teklad custom diet TD.210230 (5% Pectin, 5% Inulin with 61 kcal% of fat) was used. For the LP diet, Teklad custom diet TD.160634 (6 kcal% of protein) was used. Fecal pellets were collected from individual mice on days 10,18, 35, 43, 57, 65, and 76 in a sterile hood, transferred to 1.5 mL tubes, and placed on dry ice through the duration of sampling. Following sample collection, tubes were stored at –80° C.

## METHOD DETAILS

### SAMPL-seq analyses

Fecal pellets were recovered from –80° C storage and transferred to 1.5 mL tubes containing methacarn (60% methanol, 30% chloroform, 10% acetic acid). After 24 hrs of fixation, samples were washed with 70% ethanol and processed following the SAMPL-seq protocol^31^. After fracturing and barcoding, 20–40-micron particles were isolated by size-exclusion filtering for sequencing. For each mouse, two technical replicates of approximately 20,000 particles were used for sequencing. Samples were sequenced from single-ends for 152 cycles on an Illumina NextSeq550 using a High-output reagent Kit (Illumina 20024907).

### Bioinformatics and data pre-processing

#### Sequence filtering and 16S rDNA amplicon analysis

Raw sequencing reads were first filtered using USEARCH 11.0.667^82^, with a cutoff of less than 1 expected error and a minimum length of 150bp. Filtered reads were then demultiplexed using ULTRAPLEX^84^. Specifically, ULTRAPLEX was used to identify and extract barcodes from each read, using a custom barcode mapping, and then concatenate extracted barcodes to the fastq header for each read. After barcode extraction, 16S rRNA amplicon primers were removed from reads using SeqKit^85^. The resulting reads correspond to a 69bp 16S rRNA V4 region. All samples were then pooled together for denoising using UNOISE3^95^ and reads were mapped to zOTUs using VSEARCH^83^.

Barcodes were then subjected to error correction using the DNABarcodes package^86^ in R using a custom script available on GitHub (https://github.com/gerberlab/MCSPACE). Our barcode set allows for error correction of 1 base error, so barcodes with Hamming Distance larger than 1 were considered uncorrectable and removed. ∼96% of all barcode sequences were either correct or correctable.

#### Filtering criteria

SAMPL-seq data were pre-processed before performing inference with MCSPACE. An upper read threshold of 10000 was chosen to remove particles with large amplification, potentially caused by barcode clashes or experimental artifacts. To establish filtering thresholds for a minimum number of reads per particle and a minimum OTU abundance across all particles, threshold values were varied and the resulting numbers of particles and OTUs retained were plotted for each threshold (**Figure S1**). Final threshold values were then chosen by picking an “elbow” for each plot. Particles with numbers of reads below 250 or above 10000 were first removed. The 250 read threshold corresponded to the 95^th^ percentile in human data and the 88^th^ percentile in mouse data, while the 10000 read threshold corresponded to the 99.8^th^ percentile in human data and the 99.4^th^ percentile in mouse data. OTU relative abundances were then computed by aggregating over their abundances in the remaining particles. OTUs occurring above 0.005 relative abundance at any timepoint in human data and OTUs occurring above 0.005 relative abundance in at least 2 subjects at any timepoint in mouse data were retained, to remove OTUs that could not be reliably quantified in samples.

#### Taxonomic assignments

For the mouse dataset, taxonomy was assigned to OTUs using the RDP classifier^87^. In order to obtain better taxonomic resolution, a phylogeny-aware taxonomic identification was used for the human dataset, as in Kozlov et al^96^, using the software gappa^88^. Gappa requires phylogenetic information that is not as reliable for the mouse gut microbiome and was therefore not applied to the mouse dataset.

#### Phylogenetic placement

Phylogenetic placement was used to visualize assemblages on a phylogenetic tree. A reference phylogenetic tree constructed from full length 16S rRNA sequences was taken from Maringanti et al^97^ (https://github.com/gerberlab/mditre/tree/master/mditre/tutorials/pplacer_files). Placement of OTUs on the tree was then performed by running the software pplacer^89^ with default settings.

### MCSPACE software

MCSPACE was implemented in Python 3.10 using the PyTorch, Numpy, Scipy, Matplotlib, and Seaborn packages^90–94^. The software is publicly available as an open-source package on GitHub (https://github.com/gerberlab/MCSPACE). The input to MCSPACE consists of three csv files: (1) a file giving a table of counts for OTUs for each barcoded particle, for each sample, (2) a file giving a taxonomic label for each taxon, and (3) a file giving information on which timepoints correspond to perturbations. The software outputs inference results in two files: (a) a PyTorch file that contains the inferred model, and (b) a Python pickle file that contains summary quantities of the model posterior. The software also provides tools to visualize and interpret model results including generating pdf files of graphs for: learned spatial assemblages, assemblage proportions, and Bayes factors for significantly perturbed assemblages. See online software documentation (https://github.com/gerberlab/MCSPACE/tree/main/mcspace/docs) and tutorials (https://github.com/gerberlab/MCSPACE/tree/main/mcspace/tutorials) for complete information.

### MCSPACE model

MCSPACE uses a sparse Bayesian mixture model to discover, from co-localization count data, latent groups of spatially co-localized microbes (spatial assemblages) as well as changes in assemblage proportions across time and effects due to perturbations. Assume we have *O* OTUs, *S* subjects (biological replicates), *T* time points, and *L_sti_* particles at time-point *t*_i_ in subject *s*. The generative process for the mixture model is then as follows:

1. For each subject *s* and assemblage *k*, sample an initial untransformed mixture weight *x_ks_*(0) ∼ Normal(0,1)
2. For each subject *s*, sample the process variance 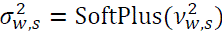, where 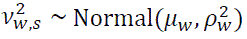
3. For each perturbation *p* and assemblage *k*:
  a. Sample perturbation indicators *c_kp_* ∼ Bernoulli(π_c_) for each assemblage *k*
  b. Sample perturbation magnitudes 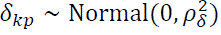 for each assemblage *k*
4. For each subsequent time point *t*_i_, assemblage *k*, and subject *s*:
  a. Compute perturbed untransformed weight means η_ks_(*t*_i_) (see below)
  b. Sample untransformed mixture weights 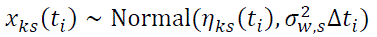
5. For each assemblage *k*, sample sparsity indicators γ_k_ ∼ Bernoulli(π_y_) and compute assemblage mixture weights 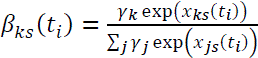
6. For each time point *t*, sample contamination mixture weights 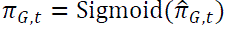, where 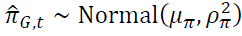
7. For each observed particle *l* in subject *s* at time *t*:
  a. Choose an assemblage assignment according to 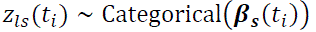
  b. For each read *j*, in particle *l*:
    i. Choose whether the read arises from the assemblage or the contamination cluster according to *a*_lj_ ∼ Bernoulli(π_G,t_)
    ii. Sample read 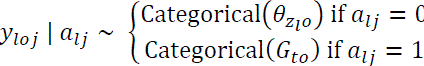

Perturbed untransformed assemblage weights η_k,s_(*t*_i_) are computed as follows:

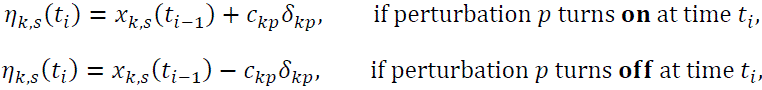

The terms *c_kp_*δ_kp_ correspond to perturbation effects. In the absence of an experimental perturbation, evolution of untransformed mixture weights is thus modelled as occurring via temporal drift.

The assemblage parameters θ_ko_, which correspond to the probability of generating reads from OTU *o* in assemblage *k*, are modeled non-stochastically and are optimized during the inference procedure. The corresponding parameters from the contamination clusters *G*_to_ are computed from data as 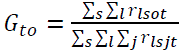, modeling the assumption that contamination arises from unencapsulated DNA that occurs with frequencies proportional to the total DNA in the sample for each OTU.

Hyperparameters were set as follows. The process variance prior location and scale were set to μ_w_ = 0.01 and 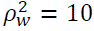, to yield a diffuse or uninformative prior. The prior probability for perturbation indicators was set to 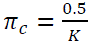, corresponding to a prior expectation of no perturbation effect on any assemblage. The prior variance for the perturbation magnitude was set to 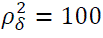, yielding a diffuse or uninformative prior. The prior location for contamination mixture weights was set to *μ_π_* = Logit(0.05) to model an expectation of 5% contamination and 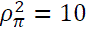 to yield a diffuse prior.

To balance the sparsity inducing prior with the influence of the data likelihood term, the prior probability π_y_ was exponentiated by a value scaled to a proportion of the totals reads in the data, e.g., pseudo-count style scaling. Specifically, we set 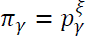, with 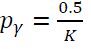, giving an expectation of less than one assemblage being present before pseudo-count scaling. We set ξ = 0.005 x #reads, where 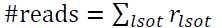, is the total number of reads in the dataset, corresponding to a prior strength of 0.5% of the dataset size.

#### Inference

Inference was performed using a Variational Autoencoder-based method^36,37^, in which parameters of the approximating distributions are estimated as functions of the data learned by a deep neural network. Specifically, inference networks were constructed that take a normalized representation 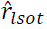 of the data as input, and output parameters for Gaussian approximating distributions, i.e. 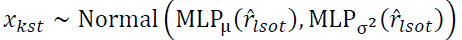. Normalized reads for each particle *l* were first passed into a fully connected 2-layer multilayer perceptron (MLP) encoder network with *O* inputs and *H* = 50 outputs, with SoftPlus activations. The outputs of the encoder were then averaged over all particles, and then passed through final linear layers that output parameters for the mean, μ, and log standard deviation, log σ, of the approximating distribution. Kullback-Leibler (KL) terms were computed analytically for all variables except for *x_kst_*, which we approximated using a Stochastic Gradient Variational Bayes estimator. Variables selecting the assemblage type, *Z*_lst_ were marginalized out for inference. A Gumbel-Softmax approach for the Bernoulli-distributed variables ℽ and *c* was used, with global parameters for each of their variational approximations^98^.

The model was implemented using Pytorch^90^ and trained end-to-end via gradient descent employing the ADAM optimizer with default parameters and initial learning rate set to 0.005. Model parameters other than θ_ko_were randomly initialized from standard Normal distributions. For initializing assemblage parameters θ_ko_, an initial fit with the *k*-means clustering algorithm was performed using the function KMeans from the sklearn.cluster package. Data input to KMeans was transformed with an Isometric Log-Ratio (ILR) transformation. For KMeans fitting, the n_clusters parameter was set equal to the MCSPACE model capacity; default values were used for all other parameters. An inverse ILR transformation was then used to map KMeans inferred cluster centers back onto the simplex, which were then used to initialize assemblage parameters θ_ko_. During inference, the sparsity prior’s exponent parameter, ξ, was gradually increased, with ξ initially set to 1 for the first 10% of training epochs, then linearly increased, reaching its final value at 90% of epochs, then held constant for the last 10% of epochs. MCSPACE was run with ten resets with different initial seeds and the model with the lowest average loss was selected.

#### Bayes Factors

Bayes factors were computed to assess the evidence of alternative models indicating the presence or absence of a perturbation effect given the data. For a given perturbation effect from perturbation *p* on assemblage *k*, Bayes factors BF_kp_ were calculated from the estimated posterior probability of the perturbation being present *q*_ckp_ and the prior probability π_c_. That is, 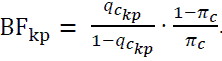.

#### Posterior summary of assemblage proportions

Posterior assemblage proportions β were summarized as follows. First, assemblages with posterior probability γ_k_ < 0.95 were removed. Latent mixture weights *x* were then sampled 1000 times from the estimated posterior distribution and passed through a softmax with discarded assemblages masked out, to get posterior samples of β^*s*^ for assemblages to be used in downstream analyses. The arithmetic mean over samples β^*s*^ was then computed to get the posterior mean of assemblage proportions.

### Comparator generative models

The scikit-learn^99^ package was used for the basic Gaussian Mixture Model. Specifically the function GaussianMixture from the sklearn.mixture package was used, with default parameters. The model was trained on the relevant input data, varying the number of components with ten runs each time with different initial seeds, and a best model was selected based on the Akaike information criterion (AIC)^100^.

For directional Gaussian mixture models, implementations provided in https://github.com/amepas/Spatial_Mbiome were used. Following Pasarkar et al^35^, each model was trained with increasing numbers of assemblages, each with ten different initial seeds, and the best fit model was selected based on the AIC. Notably, the directional GMMs failed to converge when the number of assemblages exceeded 10.

Read counts were first converted to relative abundances and then an Isometric Log-Ratio (ILR) transformation was applied as in Pasarkar et al^35^. Zeros were handled using multiplicative replacement^101^ using δ = 1/*O*^2^ for *O* taxa as in Pasarkar et al^35^.

### Benchmarking with simulated data

#### Semi-synthetic data generation

MCSPACE was run on the initial time point of the human dataset, resulting in *K* = 34 baseline assemblages. Semi-synthetic data was then generated from the baseline assemblages using the following bootstrapping-style approach:

1. Sample *K* assemblage frequencies θ and their proportions β, with replacement from the baseline assemblages. Re-normalize assemblage proportions β to sum to unity, β ←β/ ∑_k_ β_k_.
2. For each vector of assemblage frequencies θ_ko_, randomly permute OTU labels *o*.
3. Generate particle data:
  a. For each particle *l*:
    i. Sample the number of reads in particle R_l_ ∼ NegativeBinomial(*n*, *p*)
    ii. Sample an assemblage *Z*_l_ ∼ Categorical(β)
    iii. For each read *j*, in particle *l*:
      1. Choose whether the read arises from the assemblage or the contamination cluster according to *a*_lj_ ∼ Bernoulli(π_G_)
      2. Sample read 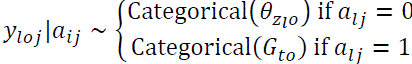

For varying the numbers of particles or reads per particle, each feature was varied individually, while the other was kept constant using default values based on what was inferred on real data. The default values used were: 1419 particles, negative binomial parameters *p* = 0.000792, *n* = 2.582442, and a contamination rate of 0.004.

MCSPACE and the basic GMM models were set to learn a maximum of 100 assemblages; as noted above, the directional GMM model failure to converge with more than 10 assemblages, so this was the maximum it was allowed to learn.

#### Comparison to pairwise analysis methods

To compare MCSPACE to methods that detect pairwise associations, a relative abundance threshold of τ = 0.005 was used to determine presence/absence of an OTU in each particle, consistent with what was used in Urtecho et al^102^. Ground-truth associations between pairs of OTUs *i* and *j* in assemblage *k* were calculated from simulation runs as *p*_ij_ = *I*(θ_ki_ > τ)*I*(θ_kj_ > τ), i.e., the association exists if and only if the abundance of both OTUs is greater than the threshold in an assemblage.

For the Fisher’s exact test and SIM9 algorithms, particle reads were first binarized. The probability of co-association with Fisher’s exact test was computed as in Sheth et al^30^. Briefly, 2 by 2 contingency tables of appearance were calculated for all pairs of OTUs and Fisher’s exact test was then used to calculate the probability of pairs occurring together more frequently than expected, assuming equiprobable occupancy at all sites. Resulting *p*-values were then adjusted via the Benjamini-Hochberg procedure^103^.

For the SIM9 approach^40^, co-association was quantified using the “sim9_single” function in the EcoSimR package^104^. On each set of particles, a binarized OTU table was subjected to a random swap, preserving OTU prevalence and particle diversities. This step was performed 25,000 times to generate a randomized community based on the original diversity of the dataset. 50 such randomized communities were generated to produce a null distribution of OTU co-localization. The location of the observed co-occurrence frequency was then determined in this null distribution and a one-tailed *p*-value was computed using a *z*-test. Resulting *p*-values were then adjusted via the Benjamini-Hochberg procedure^103^.

Co-association probabilities were computed for MCSPACE as follows. For a given posterior sample *s*, the probability that OTUs *i* and *j* co-occurred together in some particle was computed as 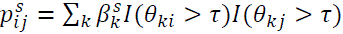. The mean over *S* = 1000 posterior samples was then computed to obtain the final probability of co-association, 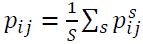.

The area under the receiver operator curve (AUC ROC) was then computed for all methods, using adjusted *p*-values for Fisher’s exact test and SIM9, and using posterior co-occurrence probabilities for MCSPACE.

#### Assemblage recovery error

The number and/or ordering of assemblages inferred by models will not in general match that of the ground truth assemblages. To account for this and provide a means to objectively compare inferred assemblages against the ground truth, we developed an algorithm that computes a metric that we term the *assemblage recovery error* (**Figure S4**). Specifically, we computed this metric as follows:

1. **Find all possible distances between inferred and true assemblages.** Specifically, we compute a matrix *D*_ij_of all pairwise distances between inferred and ground truth assemblages, using the Hellinger distance metric. The Hellinger distance is often used to quantify the distance between two probability distributions over discrete variables and is given as 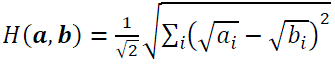, for distributions **a** and **b**.
2. **Initialize an accumulating total error.** Set the total error as *d* = 0.
3. **Match inferred and true assemblages.** While the number of rows and columns of *D*_ij_ are both greater than 1:
  a. Compute the minimum element of the distance matrix 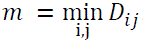, and add to the accumulated error *d* = *d* + *m*.
  b. Remove the row and column of the distance matrix *D* corresponding to the minimum element *m*.
4. **Add penalties for unmatched assemblages.** For all remaining unmatched assemblages, add a penalty value of 1.0 to the total *d*.
5. **Normalize the final error**. Normalize the error as *d* = *d*/N_max_ where N_max_ = max (N_l_, N_g_) and N_l_ is the number of learned assemblages and N_g_ is the number of ground truth assemblages.

We compute this metric over *S* = 100 posterior samples from the MCSPACE ground-truth model. For each posterior sample *s*, we only keep assemblages with 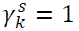, that is, those that are selected to be present in that posterior sample. The final error is summarized as the median over assemblage errors computed on each posterior sample.

For the Gaussian mixture models, inferred assemblage distributions were obtained by applying an inverse-ILR transformation on inferred cluster means, to convert them back to relative abundances.

### Cross-validation analysis with real data

For each data sample, particle data was split randomly into five test/train datasets, holding out 20% of particles in each fold. For each held-out particle, reads were down-sampled by sampling 50% of the total reads from the particle. Each model was then tasked with predicting the held out reads for each down-sampled particle.

For each cross-validated fold, MCSPACE was run with a capacity of *K*=100 assemblages. The GMM was run for a range of 2 to 100 assemblages and the best model was selected using the AIC, as described above. As noted above, the directional GMM’s failed to converge above 10 assemblages and were therefore only run up to 10 assemblages.

For GMMs, an assemblage assignment was obtained for each down-sampled particle using maximum likelihood. Predicted reads 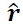 were then obtained from the assigned assemblage by first applying an inverse-ILR transformation on the inferred cluster to get relative abundances, and then multiplying the relative abundances by the number of reads of the original particle.

Each held-out particle ***r*** was then compared to the predicted reads 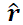 for that particle using the cosine distance: 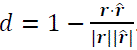.

For MCSPACE, for each held out particle ***r***, 1000 posterior samples *s* were obtained for predicted reads 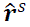, and the cosine distance was taken for each posterior sample, 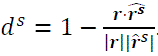. The median was then taken over posterior samples of the cosine distance *d*^s^, for each particle.

### Analysis of human and mouse datasets

#### Association analysis

To visualize changes between taxa associations over time and across conditions, we computed pair-wise association scores. We define the association score as the probability of generating a pair of reads of both OTUs *i* and *j* from the same assemblage, normalized by the marginal probabilities of generating two reads containing the OTUs: 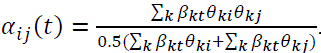

## QUANTIFICATION AND STATISTICAL ANALYSIS

### Model comparisons

For comparing model performance, statistical significance was assessed with the Wilcoxon rank sum test followed by Benjamini-Hochberg correction for multiple hypothesis testing. On semi-synthetic data, for each condition, each model was run on (*n*=10) simulated data replicates. For cross-validated performance on real data, distributions of cosine distance were compared for all particles remaining after quality filtering (*n*=7055 in human data and *n*=56848 in mouse data).

### Supplementary Tables

**Supplementary Table 1:** Composition of diets used in mouse study, in g/Kg.

|  | STANDARD | HIGH FAT | HIGH FAT, HIGH FIBER | LOW PROTEIN |
| --- | --- | --- | --- | --- |
| CASEIN | 210 | 265 | 265 | 69 |
| L-CYSTINE | 3 | 4 | 4 | 0 |
| DL-METHIONINE | 0 | 0 | 0 | 0.9 |
| LARD | 20 | 310 | 310 | 0 |
| SOYBEAN OIL | 20 | 30 | 30 | 0 |
| CORN OIL | 0 | 0 | 0 | 53.9 |
| CORN STARCH | 465 | 0 | 0 | 200 |
| MALTODEXTRIN | 100 | 160 | 160 | 0 |
| SUCROSE | 90 | 90 | 55.5 | 571.7 |
| CELLULOSE | 37.15 | 65.5 | 0 | 57.82 |
| INULIN | 0 | 0 | 50 | 0 |
| PECTIN | 0 | 0 | 50 | 0 |

### Supplementary Figures

**Supplementary Figure 1:**
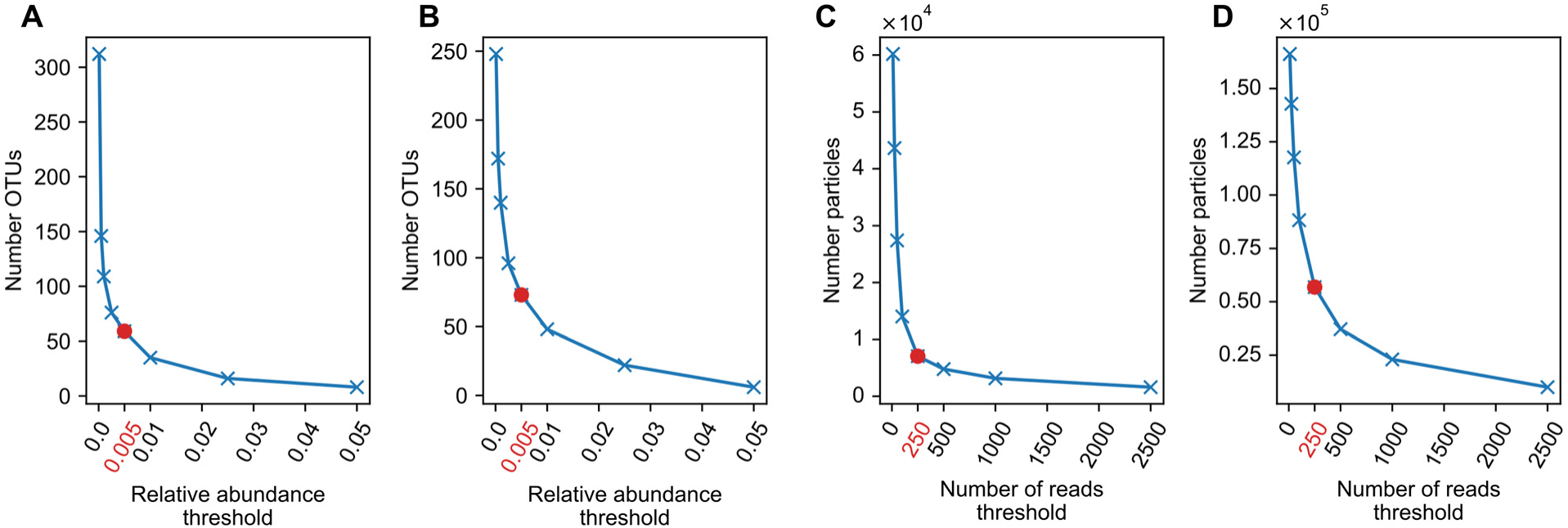
SAMPL-seq data pre-processing filtering curves. Thresholds for minimum reads per particle and OTU abundance across particles were varied, with the number of retained particles and OTUs plotted against each threshold. **(A, B)** Number of OTUs remaining versus minimum relative abundance thresholds for human **(A)** and mouse **(B)** datasets. **(C, D)** Number of particles remaining versus threshold for minimum number of reads per particle for human **(C)** and mouse **(D)** datasets. Red circles and corresponding red value on x-axis correspond to estimates of “elbows” (0.005 minimum abundance and 250 minimum reads per particle), which were then used for final data filtering.

**Supplementary Figure 2:**
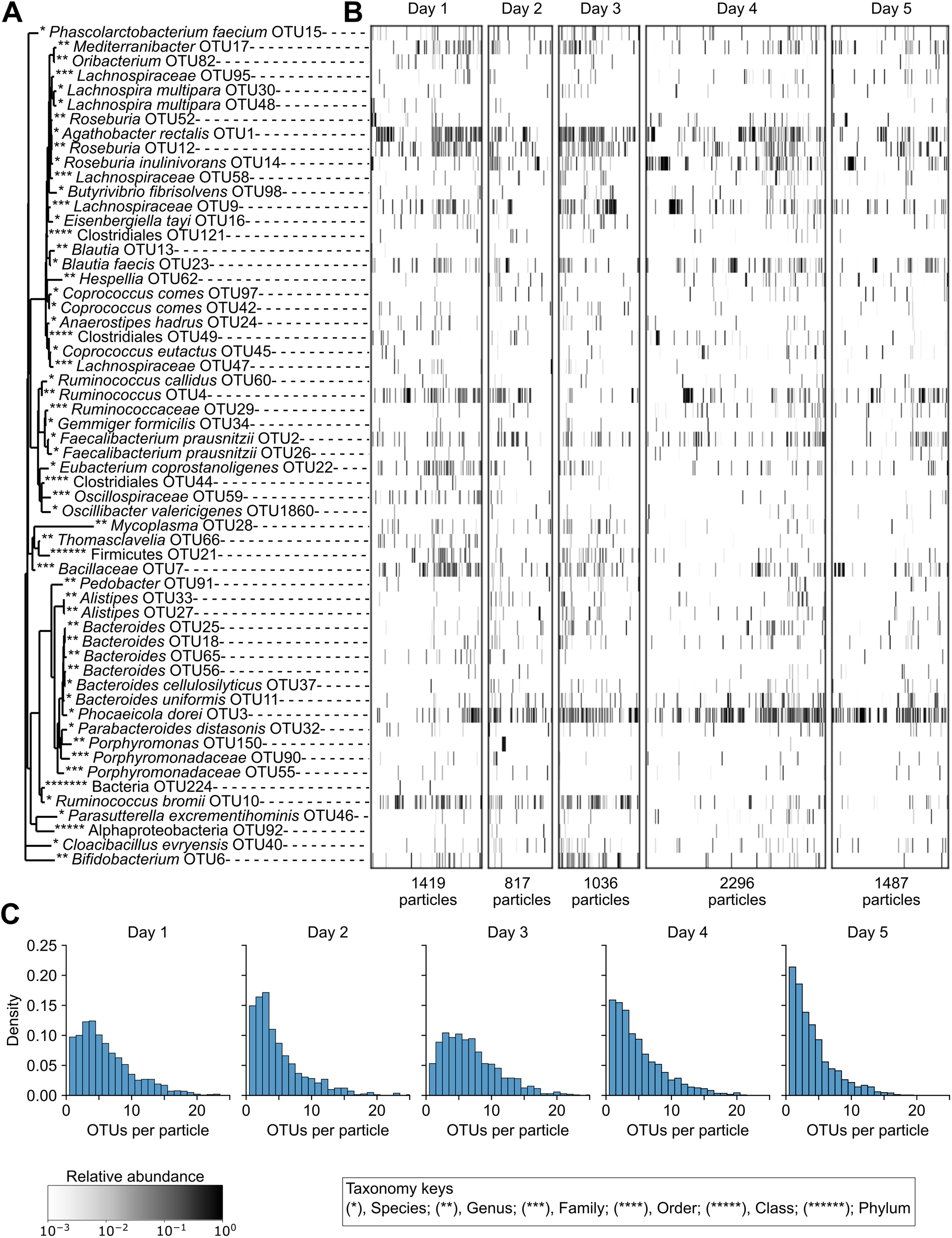
Summary of longitudinal human SAMPL-seq spatial co-localization dataset. Visualization of filtered data from a longitudinal study of gut microbiome spatial co-localization in a healthy human participant (collected daily for five days, *n*=5 fecal samples total). **(A)** Phylogenetic tree of Operational Taxonomic Units (OTUs) present in particles. **(B)** Clustered heatmap of particles, showing the relative abundance of taxa in particles over each of the five consecutive days. **(C)** Density plots of OTUs per particle.

**Supplementary Figure 3:**
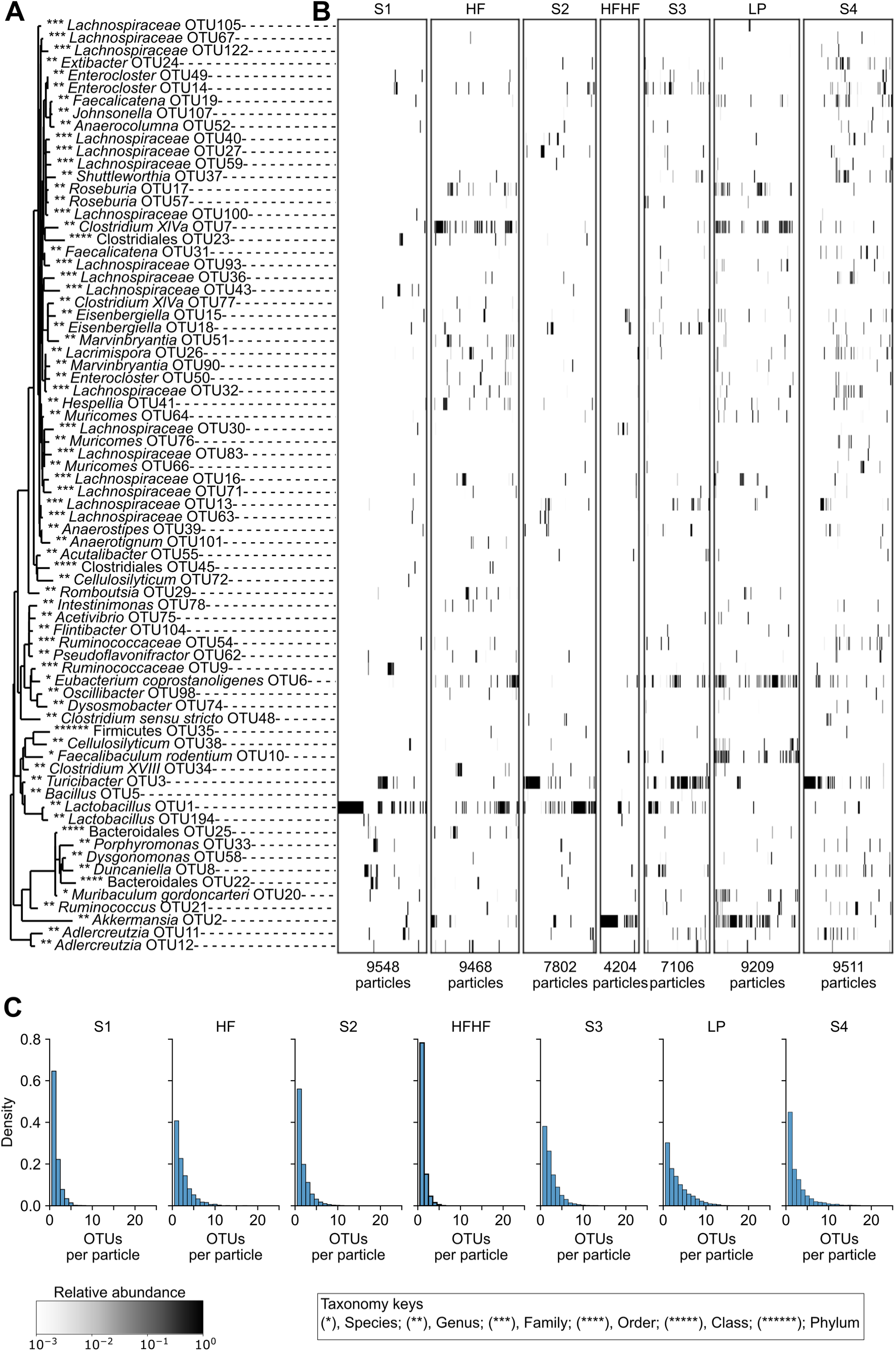
Summary of new murine longitudinal SAMPL-seq spatial co-localization dataset with multiple dietary perturbations. Visualization of filtered data from longitudinal study of gut microbiome spatial co-localization in 3 mice subjected to defined dietary perturbations (*n*=21 total fecal samples). HF = high fat; HFHF = high fat, high fiber; LP = low protein; S1-4 = standard diet 1-4. **(A)** Phylogenetic tree of OTUs present in particles. **(B)** Hierarchically clustered heatmap of particles, pooled over the three mice, showing the relative abundance of taxa in particles in each of the diets. **(C)** Density plots of OTUs per particle distributions, pooled over the three mice, for each diet.

**Supplementary Figure 4:**
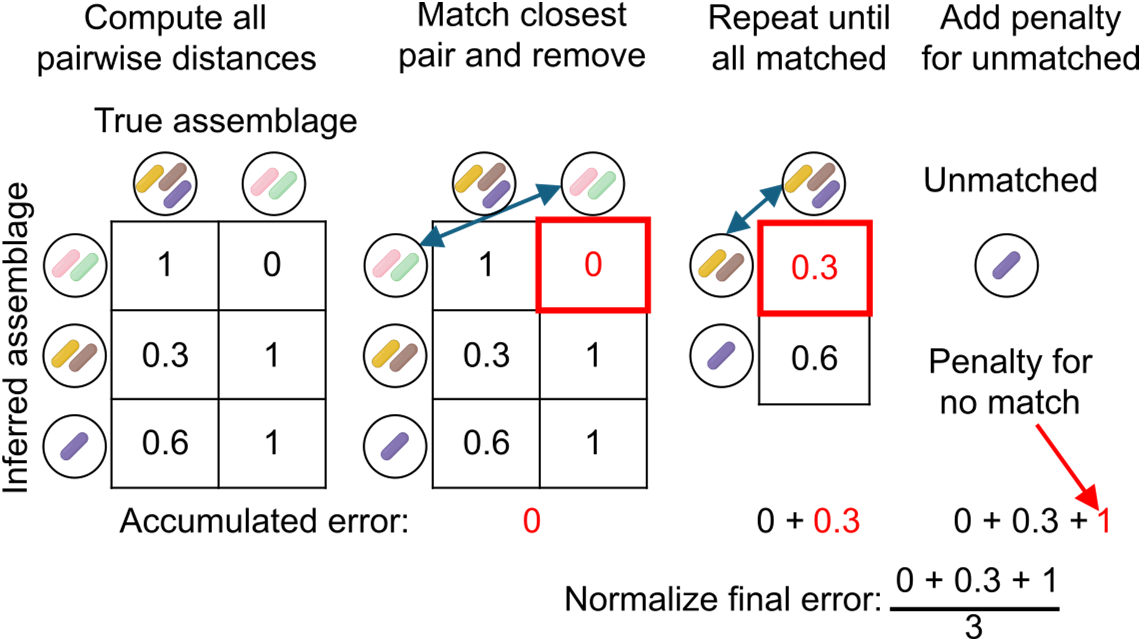
**Schematic of assemblage recovery error metric calculation**. Our metric accounts for differences in both the number and order of recovered assemblages, which may differ from that of ground truth. To compute the metric, a greedy algorithm matches inferred and true assemblages by iteratively pairing the closest matches based on pairwise distances, then accumulating errors for each pair. Unmatched assemblages incur a penalty, and the total error is normalized by the larger of the inferred or true assemblage count. Full algorithmic details are provided in the **Methods** section.

**Supplementary Figure 5:**
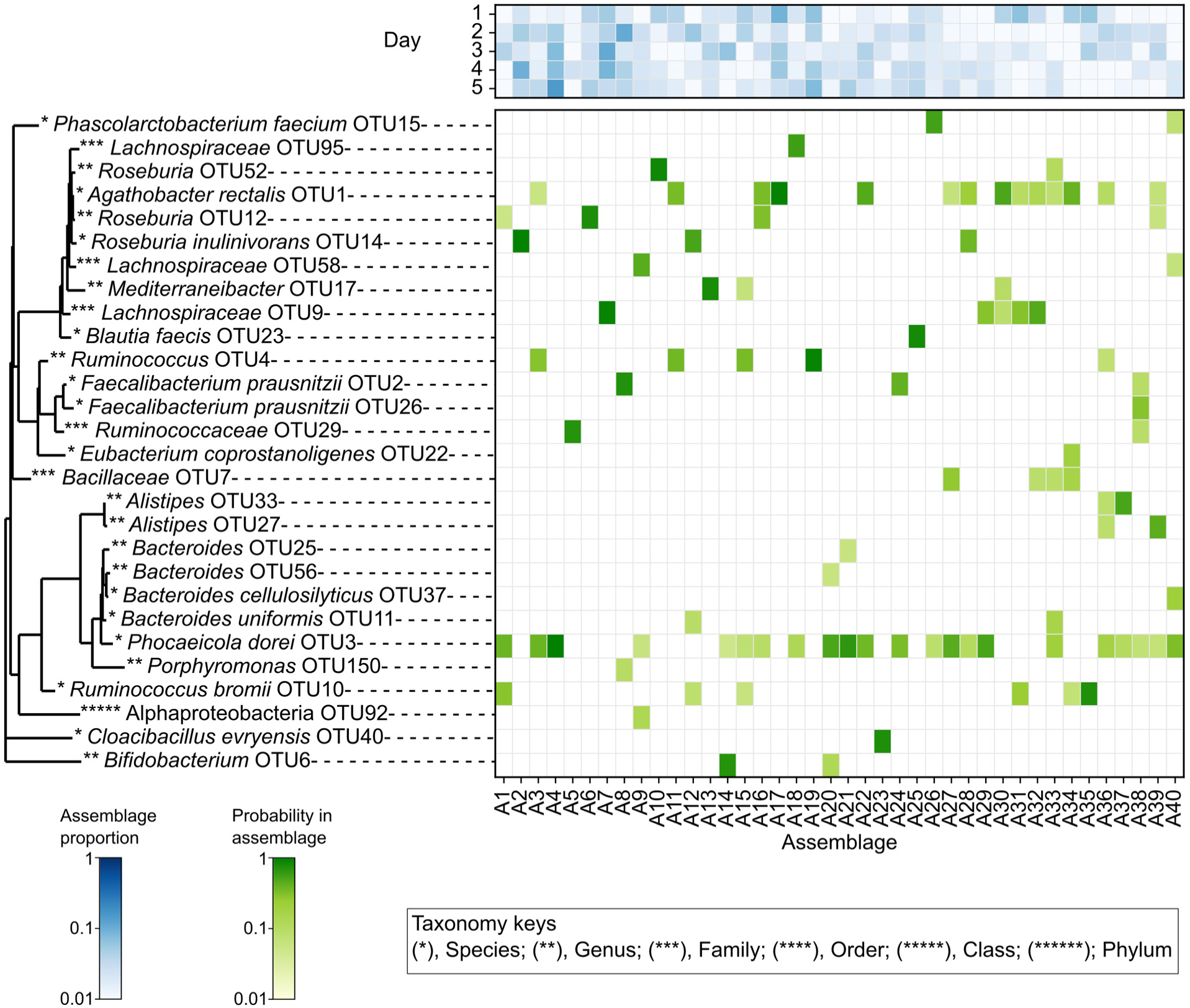
MCSPACE identified spatial assemblages among taxa and temporal changes in assemblage proportions in the human gut microbiome from a longitudinal SAMPL-seq dataset. MCSPACE identified 58 OTUs assorting into 40 spatial assemblages, with assemblage abundances tracked over time. A phylogenetic tree of OTUs present in the dataset is displayed on the left. Heatmaps show assemblage proportions over the five consecutive days in the study (above), and OTU frequencies in inferred spatial assemblages (below) with OTU assemblage frequencies ≥0.05 shown.

**Supplementary Figure 6:**
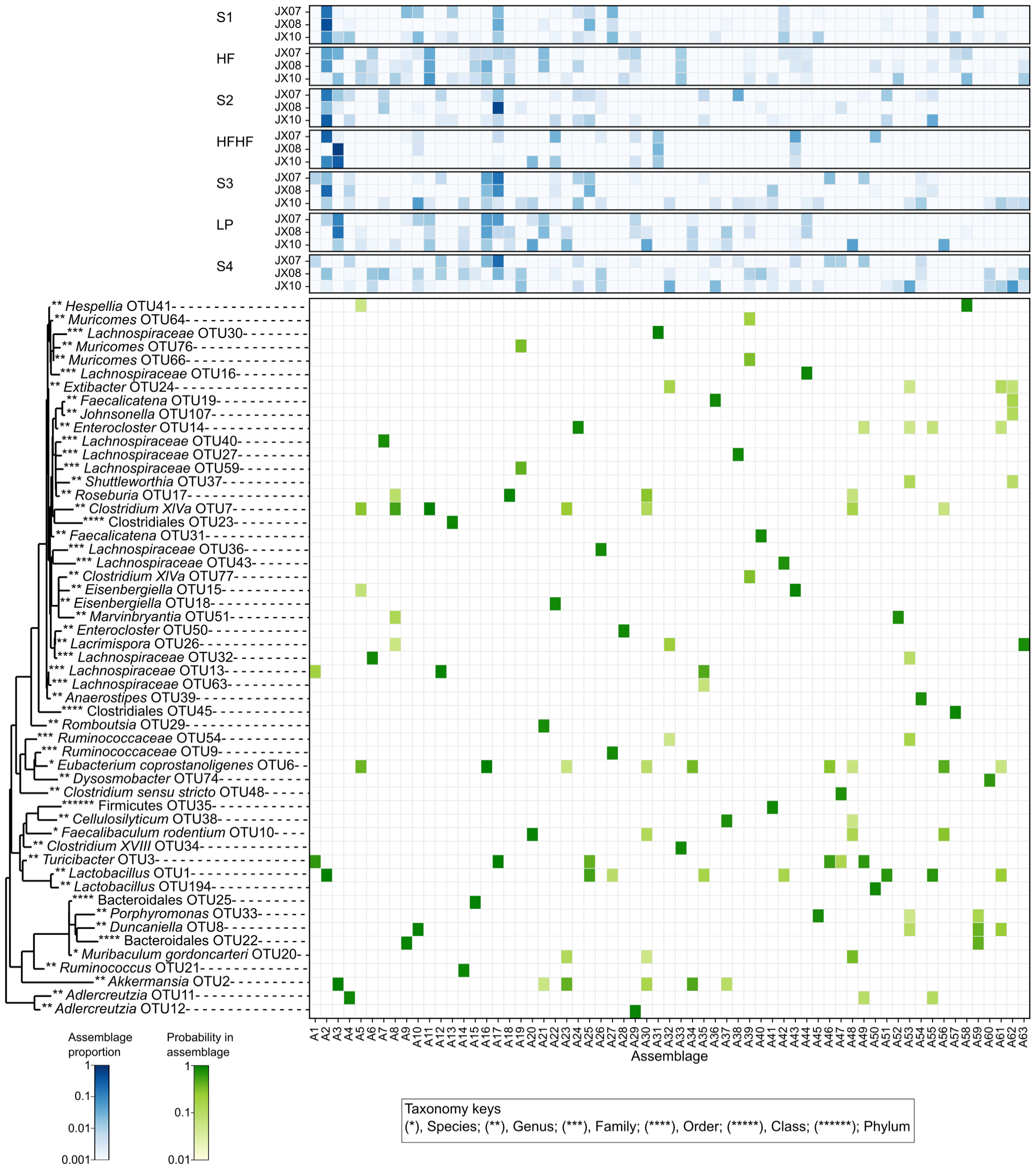
MCSPACE identified spatial assemblages among taxa and changes in assemblage proportions in the murine gut microbiome from a SAMPL-seq dataset investigating multiple dietary perturbations. MCSPACE identified 74 OTUs assorting into 63 spatial assemblages, with assemblage abundances tracked over time for three biological replicates. A phylogenetic tree of OTUs present in the dataset is shown on the left. Heatmaps show assemblage proportions (above) inferred for each biological replicate over the seven dietary intervals (HF = high fat; HFHF = high fat, high fiber; LP = low protein; S1-4 = standard diet 1-4), and OTU frequencies in inferred spatial assemblages (below). OTUs with assemblage frequencies *≥*0.05 are shown.

**Supplementary Figure 7:**
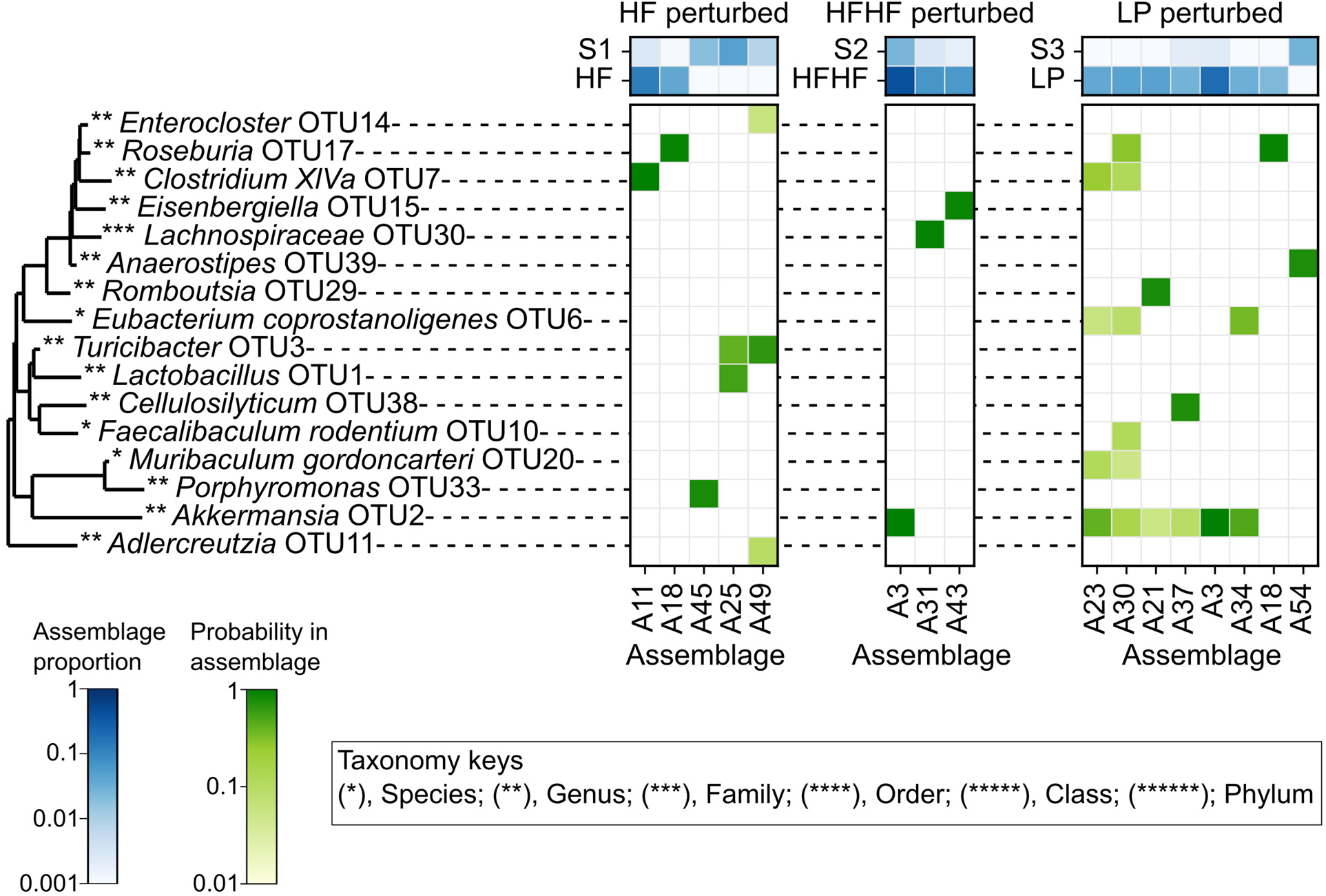
MCSPACE analysis of the murine data revealed diet-induced spatial shifts in the gut microbiome. Assemblages with strong evidence of an effect from one or more perturbations (Bayes Factors > 10) are shown for each dietary perturbation. HF = high fat; HFHF = high fat, high fiber; LP = low protein; S1-3 = standard diet 1-3. Phylogenetic tree of OTUs present with frequency >5% in any of the perturbed assemblages is shown on the left. Heatmaps show assemblage proportions before and after perturbation for each dietary perturbation (above) and OTU frequencies in perturbed spatial assemblages (below).

